# Hubs of long-distance co-alteration in brain pathology

**DOI:** 10.1101/846642

**Authors:** Franco Cauda, Lorenzo Mancuso, Andrea Nani, Jordi Manuello, Donato Liloia, Gabriele Gelmini, Linda Ficco, Enrico Premi, Sergio Duca, Tommaso Costa

## Abstract

The exact mechanisms at the root of pathologic anatomical covariances are still unknown. It is nonetheless becoming clearer that the impact of brain diseases is more convincingly represented in terms of *co-*alterations rather than in terms of localization of alterations. According to this view, neurological and psychiatric conditions might be seen as whole-brain patterns of modifications. In this context, the physical distance between two co-altered areas may provide insightful information about how pathology develops across the brain, assuming that long-range co-alterations might be relevant features of pathological networks. To demonstrate this hypothesis, we calculated the probability of co-alteration between brain areas across a large database of voxel-based morphometry studies of psychiatric and neurological disorders, and we investigated the physical (Euclidean) distance of the edges of the resulting network. Such analysis produced a series of observations relevant for the understanding of pathology, which range from unanticipated results to the recognition of regions of well-known functional and clinical relevance. For instance, it emphasizes the importance of the anterior and dorsal prefrontal cortices in the distribution of the disease-related alterations, as well as a specular asymmetry of gray matter decreases and increases between the hemispheres. Also, the analyses of schizophrenia and Alzheimer’s disease show that long-distance co-alterations are able to identify areas involved in pathology and symptomatology. Moreover, the good concordance between the measure of the mean physical distance and that of functional degree centrality suggests that co-alterations and connectivity are intimately related. These findings highlight the importance of analyzing the physical distance in pathology, as the areas characterized by a long mean distance may be considered as hubs with a crucial role in the systems of alterations induced by brain diseases.

## 1. Introduction

Converging evidence is revealing that the impact of diseases on brain structure is better appreciated not as the simple spatial distribution of lesions but as a system of interrelated alterations affecting networks (Evans, 2013). This makes sense in neurodegenerative diseases, where misfolded proteins frequently spread from one area to another in a prion-like fashion (Goedert et al., 2017; Guest et al., 2011). This mechanism has been put forward to explain the development of anatomical alterations observed in such diseases in terms of connectivity pathways, along which pathological proteins (proteinopathy) or other toxic agents can propagate (Mandelli et al., 2016; Manuello et al., 2018; Saxena and Caroni, 2011; Seeley et al., 2009; Warren et al., 2013, 2012). However, the network-like account of co-alterations seems to provide insights also in clinical conditions that do not have a neurodegenerative origin, such as schizophrenia, autism, obsessive-compulsive disorder, depression, and chronic pain (Cauda et al., 2018a, 2017, 2014; Tatu et al., 2018; Wheeler et al., 2017, 2015). Furthermore, transdiagnostic meta-analyses merging data of studies about psychiatric and neurologic diseases support the following ideas: i) the most affected areas of the brain correspond to the hubs of the functional and structural connectomes (Crossley et al., 2014), and ii) the distribution and development of co-alterations are mainly explained by functional and structural connectivity constraints (Cauda et al., 2018b). Therefore, the anatomical substrate of brain disorders might be better accounted for by patters of *co-alterations* than by the simple localization of a series of unrelated alterations.

A relevant feature of brain organization is the physical distance between interconnected areas. The small-world networks of the human connectome are composed by several clusters of short-range connectivity, linked together by long-range connections (Watts *et al*., 1998; Sporns and Zwi, 2004; Achard *et al*., 2006). Within an evolutionary perspective, the brain evolved to match a trade-off between minimizing wiring cost and allowing a fast, efficient and resilient information flow (Bullmore and Sporns, 2012; Laughlin and Sejnowski, 2003). Nodes of long-range connectivity are to be considered central hubs of the connectome, as they tend to be connected with many other nodes, which makes them crucial in the interplay between segregation and integration that shapes brain structure and function (Alexander-Bloch et al., 2013; Bullmore and Sporns, 2012; Liao et al., 2013). This organization is likely to be associated with the distribution patterns of neuropathological alterations, as brain hubs with high centrality are also the regions more likely to be affected by pathology (Crossley et al., 2014). Interestingly, both functional (Alexander-Bloch et al., 2013) and anatomical covariance (Bassett *et al*., 2008) networks (Bassett et al., 2008) of diseases such as schizophrenia, have been associated with altered values of physical distance. Higher-order associative brain regions, which are more prone to be targeted by diseases (Crossley et al., 2014), are characterized by a long physical distance and a high centrality (Sepulcre et al., 2010). So, if connectivity influences the development of pathology (Cauda et al., 2018b; He et al., 2007; Mandelli et al., 2016; Seeley et al., 2009; Zhou et al., 2012), the spatial distribution of the physical distance of co-alterations might provide an insightful indication of how pathology is distributed across the brain. It would be also interesting to compare such information with a measure of centrality from a normative connectome, so as to test if there is a correlation between abnormal distance and connectivity. This would help clarify the relationship between co-alteration and connectivity, as well as understand better the complex systems of alterations in action on the brain. In the present study we assessed the mean physical distance of co-alterations and its relationship with functional degree centrality (DC) in a meta-analytic, transdiagnostic way, so as to identify the cerebral areas that are more involved in long-range systems of pathological modifications. We obtained from the BrainMap database both voxel-based morphometry (VBM) and activation data to be used in a meta-analytic technique based on the Patel’s κ (Cauda et al., 2018a; Mancuso et al., 2019; Patel et al., 2006). This transdiagnostic approach was motivated by the assumption that the mechanisms underlying the phenomenon of co-alteration seem to be related to normative brain connectivity (Buckholtz and Meyer-Lindenberg, 2012; Cauda et al., 2018b). We therefore constructed networks of pathologic co-alterations of gray matter (GM) decreases or increases and calculated the mean physical (Euclidean) distance for each brain region. Then we compared the map of physical distance with a map of meta-analytic degree centrality of co-activations, so as to see whether or not the most connected areas of the functional healthy brain are also those with the longest co-alterations. Moreover, we investigated the transdiagnostic variability of each voxel and network, which allowed to assess the convergence of the most important pathologies in respect to their distance of co-alteration. Finally, taking schizophrenia and Alzheimer’s disease as representative examples of a single-pathology approach for psychiatric and neurologic diseases, we observed their maps of mean physical distance of co-alterations. This investigation showed that the anatomical distribution of the mean physical distance can provide an insightful index of the pathologic spread of single diseases.

## 2. Materials and methods

### 2.1 Collection of data

In the present study the Cochrane Collaboration definition of meta-analysis (Green et al., 2008) was adopted and the selection of articles was conducted referring to the “PRISMA statement” international guidelines (Liberati et al., 2009; Moher et al., 2009). Neuroimaging experiments eligible for the analysis were retrieved from BrainMap (http://brainmap.org/) (Fox et al., 2005; Fox and Lancaster, 2002; Laird et al., 2005; Vanasse et al., 2018). BrainMap is an open access online database constituted by over 15000 neuroimaging experiments and 120000 locations in stereotaxic brain space. The database has two sections for VBM and functional data sets.

The VBM BrainMap section was queried (November 2017) with the following algorithms:

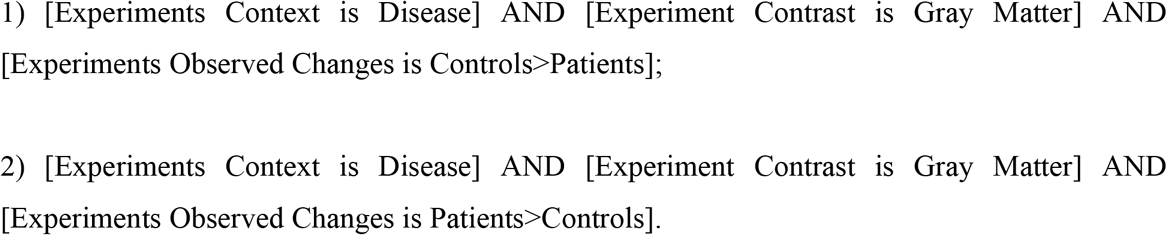

The first step consisted in the codification of the VBM data set following the ICD-10 classification (World Health Organisation, 1992). Subsequently, full-text articles were analyzed in order to verify that they conformed with the following criteria: a) being an original work published in a peer-reviewed English language journal; b) performing a whole brain VBM analysis technique; c) including a matched comparison between a pathological group and a group of healthy subjects; d) reporting GM decrease/increase changes in the pathological sample; e) adopting a specified VBM analysis; f) referring to a specific stereotaxic space (e.g. Montreal Neurological Institute space or Talairach/Tournoux atlas) as regards GM increase/decrease changes. On the basis of these criteria we deemed eligible 912 experiments for GM decreases and 350 for GM increases. Descriptive information of interest was extracted from each full-text article fulfilling the abovementioned criteria. As a further specification, all the experiments not coded with F (i.e. mental, behavioral and neurodevelopmental disorders) or G (i.e. diseases of the nervous system) labels were excluded from the analysis. Moreover, studies related to codes that could not be considered as primary brain disorders (i.e. F10: Alcohol related disorders; F15: Other stimulant related disorders; F28: Other psychotic disorder not due to a substance or known physiological condition; F91: Conduct disorders; G11: Hereditary ataxia; G43: Migraine; G44: Other headache syndromes; G47: Sleep disorders; G50: Disorders of trigeminal nerve; and G71: Primary disorders of muscles) were also excluded. Articles including less than 8 subjects were excluded as well. This lower bound was chosen in accordance with the work of Scarpazza and colleagues (Scarpazza et al., 2015), whose work demonstrated that VBM experiments based on an equivalent sample should not be biased by an increased false positive rate. At the end of the selection procedure, 642 remaining experiments for the GM decreases (for 15820 subjects and 7704 foci) and 204 remaining experiments for the GM increases (for 4966 subjects and 2244 foci) were included in the analyses. We also calculated two single-disease co-alteration networks using only on the data of schizophrenia and Alzheimer’s disease (see Supplementary Fig. 1 and Supplementary Table 3 and 4).

Finally, a systematic search was conducted through the entire functional data set of BrainMap with the following query:

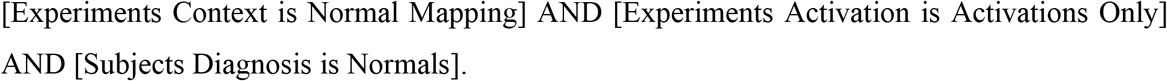

This search produced 2376 articles, for a total of 13148 experiments, 112 paradigm classes and 68152 subjects (see Supplementary Fig. 1 and Supplementary Table 6). Finally, we converted all locations reported in MNI into Talairach space using the Lancaster’s icbm2tal transform (Lancaster et al., 2007).

### 2.2 Modeled alteration maps

To obtain the alteration maps from the BrainMap foci, we adopted the anatomical likelihood estimation (ALE) framework (Eickhoff et al., 2009, 2012; Turkeltaub et al., 2012) to produce a modeled alteration (MA) map for each experiment. To build the MA maps, every focus of each experiment is taken as the central point of a 3-dimensional Gaussian distribution of probability:

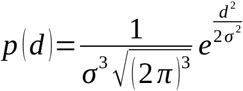

In this formula *d* represents the Euclidean distance between the voxels and each considered focus, while *e* indicates the spatial uncertainty. The standard deviation is obtained through the full-width half-maximum (FWHM), which is defined as follows:

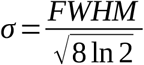

### 2.3 Co-alteration matrix calculation

The brain was partitioned on the basis of an anatomical atlas derived from the Talairach Daemon atlas with a resolution of 2 mm (Lancaster et al., 2000); a co-alteration matrix resulted from all the possible couples of brain areas. In the resulting ExN matrix, each of E row stands for an experiment, while each of N column reports a node; in the present study the matrix included 642 experiments (VBM contrasts) x 1105 nodes for the decrease condition, and 204 experiments x 1105 nodes for the increase condition. For each experiment, a given brain region was considered to be altered if the experiment MA map showed 20% or more of significant voxels within the region. The choice of this threshold is arbitrary, but it has already been proven that it do not affect the results significantly (Mancuso et al., 2019). The use of Patel’s κ index allowed us to obtain the co-alteration strength between the nodes (Patel et al., 2006); then, a probability distribution of joint alteration occurrences for every couple of nodes was created following a Bernoulli model of data generation. In other words, given two VOIs (*a* and *b*), we can determine all their conjoint states of alteration as follows: i) *a* and *b* are both altered; ii) *a* is altered and *b* is not; iii) *b* is altered and *a* is not; iv) neither *a* nor *b* are altered. Thus, frequencies of these four cases recurring in all the experiments could be described by the following probability formulas:

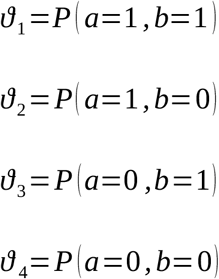

These probabilities stand for the conjoint state frequencies of a couple of VOIs (*a* and *b*) in the four possible combinations. Table 1 contains the marginal probabilities. Obtaining these probabilities allowed us to use the Patel’s *κ* index to create the correlation matrix including each couple of brain areas. Patel’s *κ* informs about the likelihood that two VOIs (*a* and *b*) are co-altered, namely, it expresses the probability that *a* and *b* are altered independently of each other. Patel’s *κ* is defined as follows:

**Table 1:** Marginal probabilities between altered and unaltered regions.

| VOI b | VOI a |  |  |  |
| --- | --- | --- | --- | --- |
|  |  | Altered | Unaltered |  |
| | Altered | $\vartheta_1$ | $\vartheta_3$ | $\vartheta_1 + \vartheta_3$ |
| | Unaltered | $\vartheta_2$ | $\vartheta_4$ | $\vartheta_2 + \vartheta_4$ |
| | | $\vartheta_1 + \vartheta_2$ | $\vartheta_3 + \vartheta_4$ | 1 |

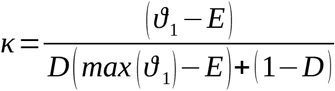

where

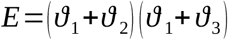

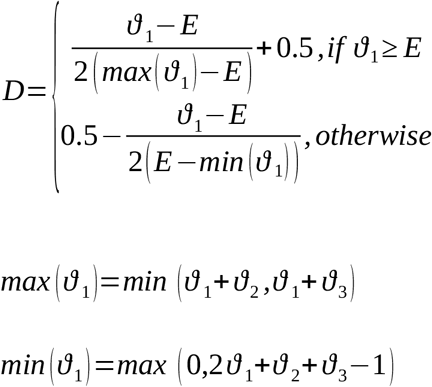

The numerator of the fraction is the difference between the likelihood that *a* and *b* are altered together and their expected joint alteration in a condition of independence, and the denominator is a weighted normalizing constant. *Min*(*ϑ*_1_) represents the maximum value of conjoint probability *p* (*a, b*), given *p* (*a*) and *p* (*b*), whereas *max* (*ϑ*_1_) represents the minimum value of *p* (*a, b*), given *p* (*a*) and *p* (*b*). Patel’s *κ* index returns values ranging from –1 and 1. A value of |*κ*| that is close to 1 characterizes a high co-alteration. The statistical significance of Patel’s *κ* is assessed by means of a simulation of a generative model of data based on the Monte Carlo’s algorithm. Specifically, the algorithm computes an estimate of *p*(*κ*|*z*) by sampling a Dirichlet distribution and determining the proportion of the samples in which *κ* >*e*, where *e* is the threshold of statistical significance. The resulting co-alteration matrix returns values that are proportional to the statistical relationship between the patterns of brain areas’ alterations taken into account.

### 2.4 Functional connectivity matrix

The meta-analytic connectivity was obtained applying the same method for the construction of the co-alteration matrix (i.e. Patel’s *κ* index calculated on each couple of brain areas) to the BrainMap functional database of activations of healthy subjects.

### 2.5 Measurement of the mean distance and calculation of the meta-analytic degree centrality

For each significant statistical association between two regions *a* and *b* in the co-alteration (or co-activation) matrix, we calculated the respective physical distance as the Euclidean distance between the centroids of *a* and *b* in the Talairach Daemon atlas. Then, for each node, the distance of its connections was averaged to obtain its mean physical distance, and such value was assigned to the whole region to obtain a map.

To calculate the degree centrality (DC) of co-activation on the functional data we employed the Brain Connectivity Toolbox algorithms (Rubinov and Sporns, 2010). Specifically, DC was defined as the number of edges incident to a node.

### 2.6 Comparison between maps

In order to evaluate the similarity between the distance of the co-alteration map and the functional DC map, the correlation between these maps was calculated. To establish the involvement of each brain area to the correlation between the two maps we applied the voxels’ contribution to correlation (VCC) technique (Mancuso et al., 2019). This *leave-one-out* procedure recursively computes the correlation between each couple of maps, subtracting a voxel at every run (the same voxel for both the maps). To create a map showing the contribution of each voxel to the correlation, the difference between the correlation value calculated in the whole maps and the correlation value obtained after the exclusion of each couple of voxels is associated to the singular voxel. Therefore, it is possible to visualize the extent to which the correlation value decreases or increases depending on the subtraction of each couple of corresponding voxels from the calculation. The normalized version of this map (transformed in order to range from –1 to 1) represents the stability of correlation and describes the contribution of each voxel to the correlation. This procedure allows to characterize voxels as positive or negative. The former contribute positively to the correlation, as their removal decreases the correlation value; the latter contribute negatively to the correlation, as their removal increases the correlation value.

### 2.7 Leave-one-pathology-out

The leave-one-pathology-out is a validation technique used to evaluate both the variability and generalizability of measurements. The procedure for the calculation of the mean physical distance described above was performed several times, each run excluding a different pathology. For all the outcomes resulting from each measurement the voxel-wise standard deviation was calculated to verify the degree of variability and consistency of the different measurements with regard to each pathology.

## 3. Results

### 3.1 Maps of the mean physical distance

The map of the mean physical distance of co-alterations related to GM decreases show higher peaks in the dorsal and anterior regions of the left prefrontal cortex (PFC), and the bilateral medial temporal lobe (MTL). Left posterior insula, left postcentral gyrus, right precentral gyrus and clusters in the bilateral temporal and occipital lobes are also involved. The map of GM increases is characterized by more extreme values compared to that of GM decreases, thus, despite the low magnitude of many voxels, certain areas reach higher values than those observed in the other map. Those areas are the bilateral pre- and postcentral gyri, the right anterior PFC, an inferior cluster in the bilateral occipital cortex, and the left medial cuneus (Fig. 1).

**Figure 1.**
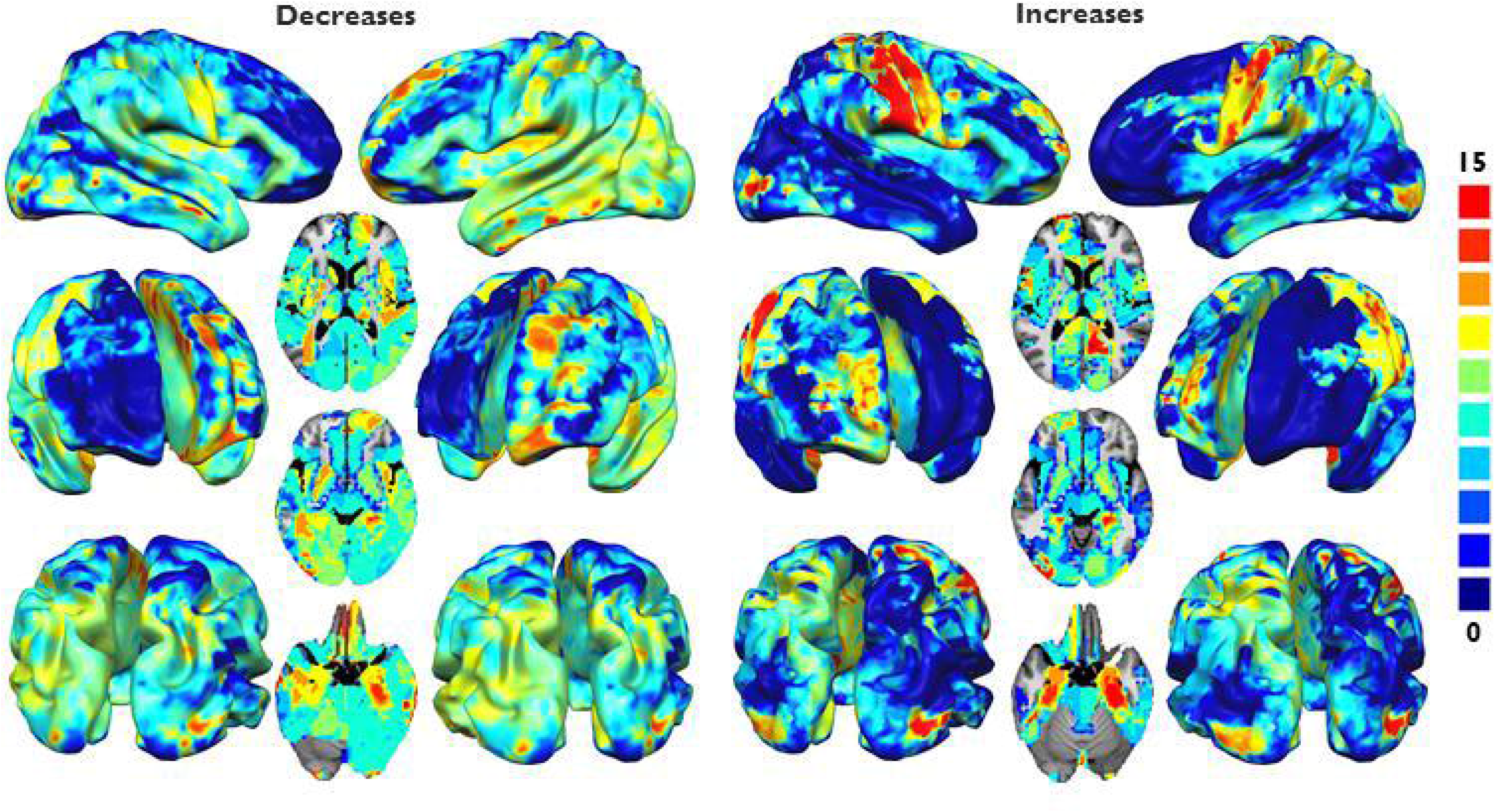
Parametric mapping of the mean distance of co-alterations, divided for decreases and increases. Higher values indicate increasing mean distance.

The sensorimotor network, the default mode network (DMN), the salience network (SN), the dorsal attention network (DAN), and the thalamus and basal ganglia are the systems where both GM increases and GM decreases show long distance co-alterations. GM decreases show longer distance co-alterations than the GM increases in the auditory network and the cerebellum (Fig. 2).

**Figure 2.**
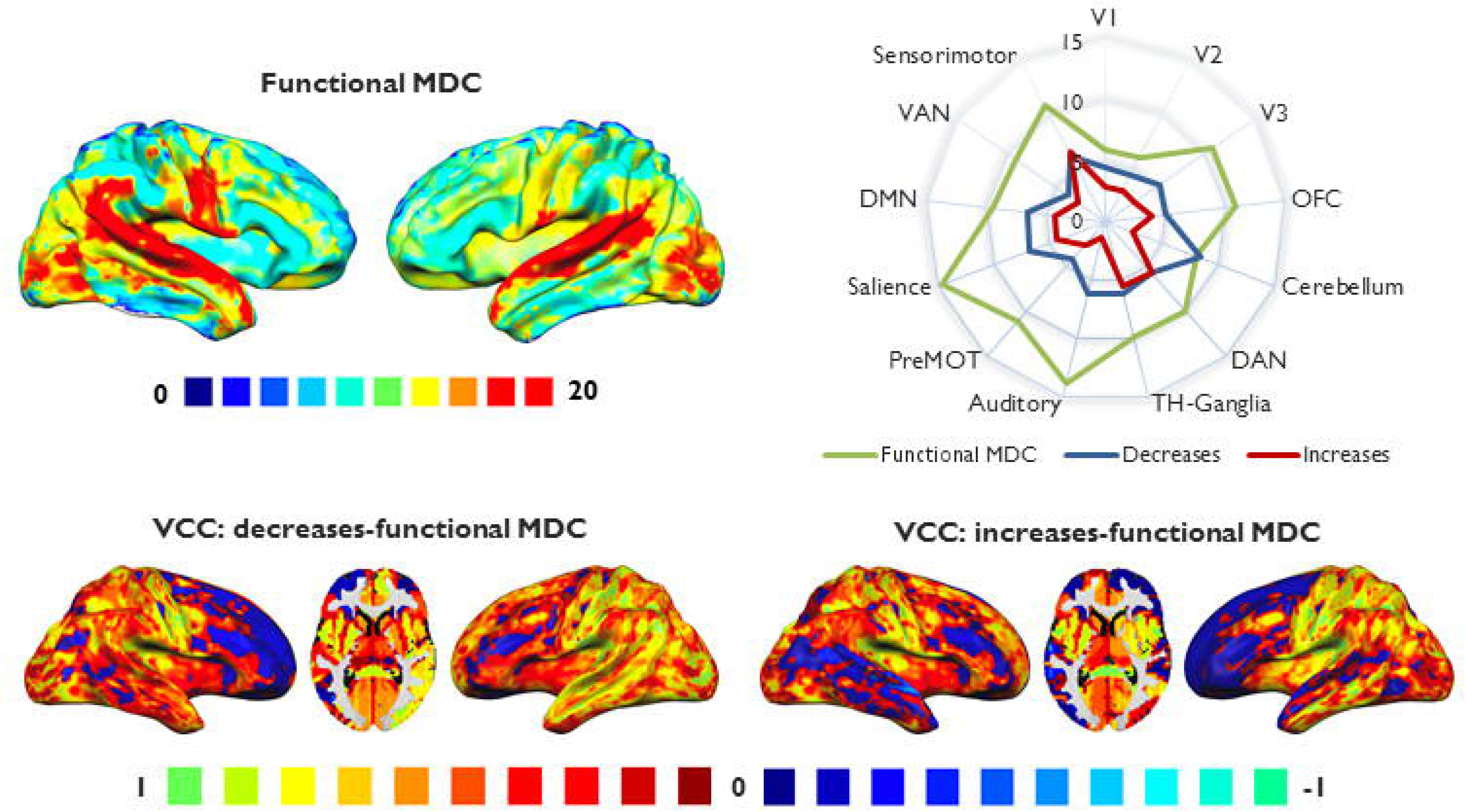
Comparison between maps of mean distance and the functional degree centrality. **Top left panel:** surface mapping of the functional meta-analytic degree centrality (functional MDC) obtained with co-activations. Higher values indicate higher degree centrality. **Top right panel:** radar chart comparing the average values of mean distance of decreases, increases and of functional meta-analytic degree centrality for each of the following networks: visual networks 1, 2 and 3 (V1, V2 and V3), orbitofrontal cortex (OFC), cerebellum, dorsal attentional network (DAN), thalamus and basal ganglia, auditory network, premotor network, salience network, default mode network (DMN), ventral attentional network (VAN), and sensorimotor network. **Bottom panel:** parametric mapping of the voxels’ contribution to correlation analysis between the functional meta-analytic degree centrality map and the mean distance co-alteration maps of decreases and increases. Positive values indicate areas of concordance between the maps, negative values indicate discordance.

### 3.2 Comparison with the map of functional degree centrality

The functional meta-analytic DC map reveals the presence of normative hubs in the bilateral superior temporal cortex, bilateral occipital cortex, right temporoparietal junction and right inferior prefrontal gyrus. The systems that present the longer distance co-alterations are the sensorimotor network, the SN and the auditory network. The correlations between the functional DC map and the maps of GM decreases and increases are *r* =0.73 and *r* =0.57, respectively. The VCC analysis reports a high concordance between the functional degree map and both the GM increases and decreases mean distance co-alteration maps in most of the voxels, except for those belonging to the right PFC and the left middle frontal gyrus (MFG) in the GM decreases map, and those belonging to the left PFC, right MFG, and bilateral temporal cortices in the GM increases map (Fig. 2).

### 3.3 Leave-one-pathology-out analysis

The regions with the highest variability across diseases are the bilateral (but especially right) inferior frontal gyrus (IFG), the bilateral insula, the bilateral temporal lobe and the bilateral occipital lobe for the map of GM decreases; the right pre- and postcentral gyri, the left MFG, the left angular gyrus and the right occipital lobe for the map of the GM increases (Fig. 3). The two maps involve different systems, especially the sensorimotor network, the DMN, the SN, the auditory network and the thalamus and basal ganglia for the GM decreases, and the sensorimotor network and V1 for the GM increases (Fig. 3).

**Figure 3.**
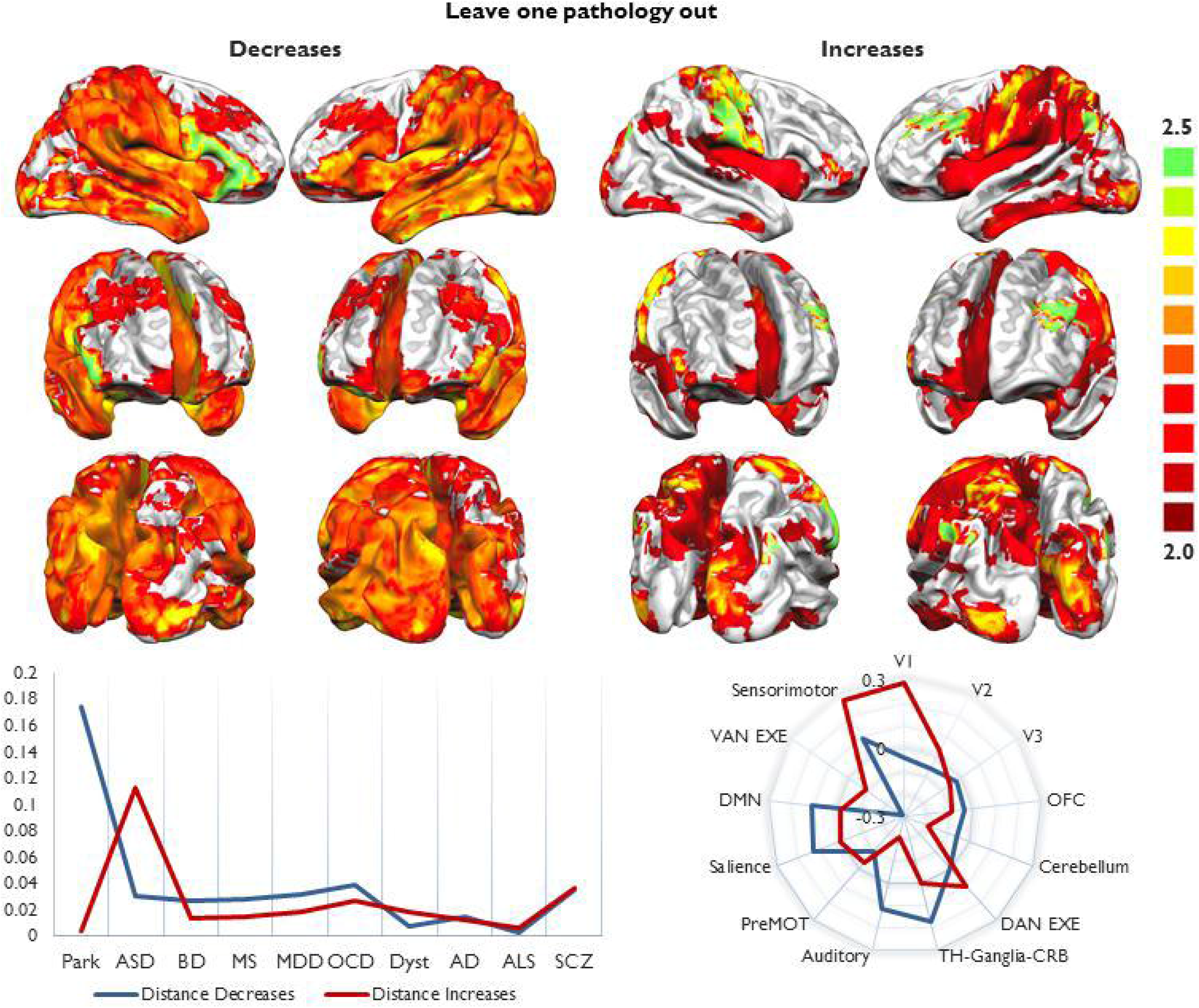
**Top panel:** surface mapping of the transdiagnostic variance of the mean distance maps of decreases and increases. Higher values indicate higher variance, that is, the voxels whose value in the mean distance map is more variable across pathologies. **Bottom left panel:** spatial variability of the changes introduced removing each pathology from the database. Higher values represent the pathologies whose changes were more uneven across resting state networks. **Bottom right panel:** values of variance of the mean distance of decreases and increases divided by resting state networks: isual networks 1, 2 and 3 (V1, V2 and V3), orbitofrontal cortex (OFC), cerebellum, dorsal attentional network (DAN), thalamus and basal ganglia, auditory network, premotor network, salience network, default mode network (DMN), ventral attentional network (VAN), and sensorimotor network.

Extracting a pathology from our database produced changes that could be unevenly distributed between the networks. For instance, the removal of Parkinson’s disease from the GM decrease dataset produced changes with the widest spatial variability, but when studies about this pathology were removed from the GM increase database the produced changes were mostly homogeneous. On the contrary, amyotrophic lateral sclerosis, Alzheimer’s disease and schizophrenia were the disorders whose removal produced the most similar impact regarding the spatial variance of changes on both the maps of GM increases and decreases (Fig. 3). These differences cannot be explained solely by the number of experiments of each pathology in the database, as disorders well represented as schizophrenia and Alzheimer’s disease had a minor impact on the results when removed compared to other less represented disorders (see Supplementary Tables 1, 2 and 5). Although analytic comparisons between disorders cannot be made, because of the uneven distribution of experiments, these findings can be considered as an indication of how different pathologies are characterized by various patterns of co-alteration distances.

### 3.4 Schizophrenia and Alzheimer’s disease

With regard to the map of GM decreases of schizophrenia (Fig. 4), the left auditory cortex shows the longest mean distance co-alterations (and, albeit less strongly, the right auditory cortex is also involved). Small clusters can be found in the right superior temporal sulcus (STS). The bilateral insula is widely involved, as well as the left IFG (especially in its posterior portion), the bilateral anterior cingulate cortex (ACC), the MTL (especially the left one), and the bilateral caudate. The map of GM increases shows long distance co-alterations mainly in the left putamen.

**Figure 4.**
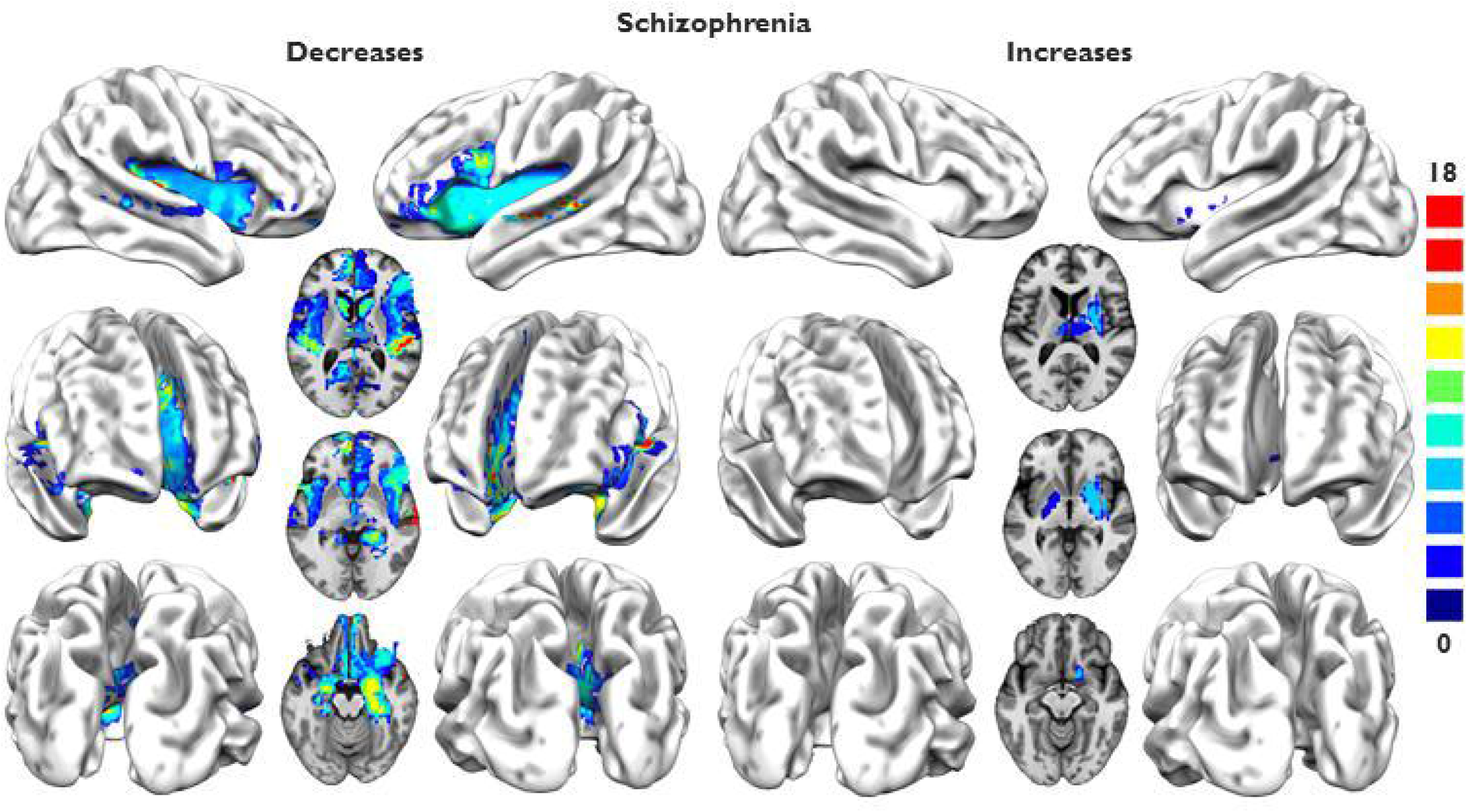
Parametric mapping of the mean distance of co-alterations in schizophrenia, divided for decreases and increases. Higher values indicate increasing mean distance.

With regard to the map of GM decreases of Alzheimer’s disease (Fig. 5), long distance co-alterations characterize the caudate (especially the left one), the MTL (especially the left one), the bilateral IFG pars orbitalis, the left orbital cortex, certain clusters in the bilateral anterior insula, and a cluster in the STS. In turn, the map of GM increases shows long distance co-alterations just in the bilateral MTL (especially the left one).

**Figure 5.**
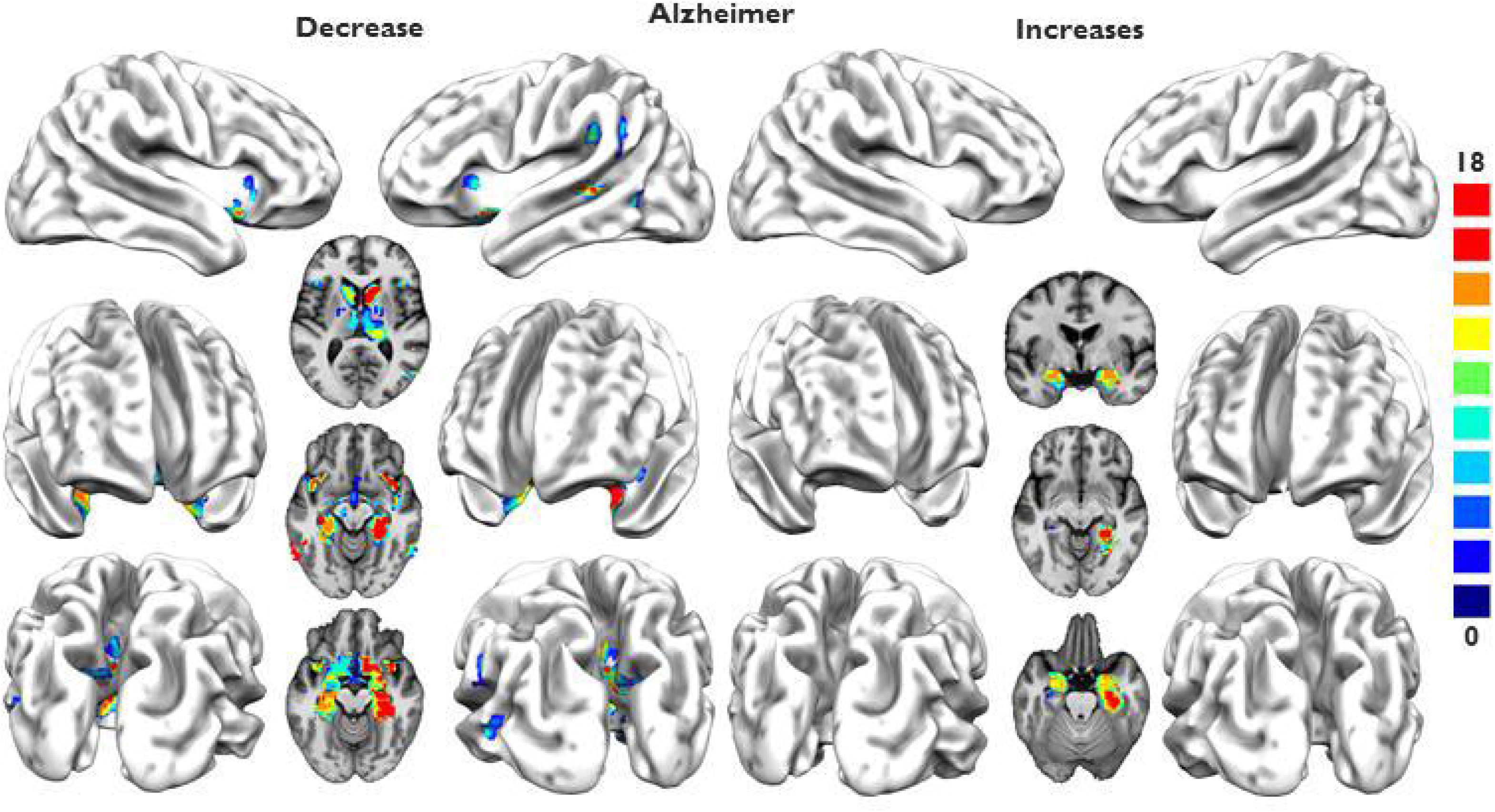
Parametric mapping of the mean distance of co-alterations in Alzheimer’s disease, divided for decreases and increases. Higher values indicate increasing mean distance.

## 4. Discussion

### 4.1 Spatial distribution of the mean distance of co-alterations

This study investigates, for the first time, the spatial distribution of the physical distance of co-alterations, highlighting that this type of measurement is able to provide insightful indications about the distribution patterns of GM alterations related to brain diseases. Findings show a spatial distribution of the mean physical distance of transdiagnostic co-alterations that can vary between areas in interesting and meaningful ways. For instance, GM decreases exhibit longer distance co-alterations in the left hemisphere compared to the right one (Fig. 1). Intriguingly, this suggests an asymmetry in the pathological involvement of the left and right hemispheres, as the former appears to show more global patterns of co-alterations, while the latter appears to be characterized by more local covariances. The left dorsal and anterior PFC tend to co-alter with distant areas. In addition, the lateral parts of PFC show low transdiagnostic variance (Fig. 3), which means that they present long-range co-alterations related to GM decreases in a wide range of pathologies. This is consistent with the low mean distance variability exhibited by the DAN (Fig. 3), as well as with the clinical observation of executive functions deficits in many diseases (Goodkind et al., 2015; McTeague et al., 2016). On the contrary, the left medial anterior PFC, and especially a part of left medial dorsal PFC, display high variance, thus suggesting that the medial PFC may be involved in long-range co-alterations in several disorders.

The MTL shows long-range co-alterations related to GM decreases. Given its involvement in memory and learning, it is likely for it to be co-altered with many other associative areas in a variety of diseases, causing symptoms of memory loss, for instance in neurodegenerative conditions such as Alzheimer’s disease, or symptoms of inappropriate memory and limbic responses that are frequently observed in the ruminations characterizing depression (Sheline et al., 2009). Significantly, the MTL is part of the DMN (Andrews-Hanna et al., 2010; Buckner et al., 2008), which is one of the most involved brain networks in long-range GM decreases (Fig. 3).

It is interesting to observe that the map of the mean physical distance of co-alterations related to the GM increases shows higher values in the right hemisphere than in the left, in spite of the left prevalence of the map related to the GM decreases. This difference is particularly noted in the dorsal and anterior lateral PFC, which is a region of significant longer co-alterations in the left hemisphere of the GM decreases’ map, while in the GM increases’ map this region presents longer co-alterations in the right hemisphere (Fig. 1). Such an asymmetry between the maps of GM increases and of GM decreases might be due to an effect of compensation (Cauda et al., 2014). In other words, while the left PFC is pathologically involved in a network of long-distance co-alterations of GM decreases the right PFC could compensate the disruption of its left homologue by being involved in a network of long-range increases. However, it cannot be excluded that both GM increases and GM decreases are primary effects of the disease itself. Future studies are needed in order to corroborate these findings and to explain the mechanism underlying this interesting asymmetry.

Other areas of long-distance co-alterations related to the map of GM increases are the pre- and postcentral gyri, particularly within the right hemisphere. Indeed, the sensorimotor network is one of the most involved systems in the map of GM increases (Fig. 3). However, these regions are also characterized by high transdiagnostic variance (Fig. 2). Overall, the sensorimotor network exhibits long-distance co-alterations related to the map of GM increases, but its involvement is not consistent across diseases; in turn, the anterior and dorsal lateral PFC are characterized by a greater transdiagnostically involvement in both the maps of GM decreases (in the right hemisphere) and of GM increases (in the left hemisphere).

### 4.2 The map of physical distance of co-alterations resembles that of functional degree centrality

The investigation of the relationship between centrality and distance) provides evidence of a good convergence between the map of co-activation DC and those of physical distance of co-alteration, especially with the one related to the GM decreases. This strongly suggests that brain functional hubs are also the regions whose mean distance of co-alteration is longer. Save for the PFC, most of the cortical areas contribute positively to the correlation between functional DC and co-alteration distance. It should be observed that the frontal voxels in the dorsal and anterior PFC displaying low convergence are not characterized by long transdiagnostic distance. On the contrary, the dorsal and anterior PFC show high convergence between co-activation centrality and co-alteration distance in the left hemisphere for the GM decreases, as well as in the right hemisphere for the GM increases, where long-distance co-alterations are found. In general, regions of long-distance co-alteration, such as the pre- and postcentral gyri and the insula, exhibit high consistency with the map of functional centrality (Fig. 2). Of note, pre- and postcentral gyri are found to be hubs of long-distance connectivity when short-rage connections are subtracted from the connectome (Esfahlani et al., 2019).

One of the fundamental issues in the study of co-alterations is to understand the mechanism that is responsible for morphological co-variations of specific sets of areas in relation to a pathological process. Our analyses provide evidence of an interesting association between the distance of co-alterations and the functional DC. As brain hubs are supposed to be preferentially targeted by pathological alterations (Buckner et al., 2009; Crossley et al., 2014; de Haan et al., 2012), the finding of a convergence between functional DC distribution and long-distance co-alteration areas suggests that brain regions which are more likely to be affected by diseases are also more likely to be co-altered with distant areas. A possible explanation might be that such regions are more susceptible to metabolic stress in virtue of their numerous connections and intense work of integration of different information (Crossley et al., 2014). Although capable of explaining the high rate of damage undergone by hubs, this model alone fails to explain our findings: it would be unclear why areas of high metabolic stress should be co-altered with other distant regions. Conversely, the transneuronal spread hypothesis (Zhou et al., 2012), which accounts for the pathologic progression with the diffusion of toxic agents such as misfolded proteins, seems to be more up to the task.

The idea that misfolded proteins are responsible for the pathological spread has found several supporting evidence in studies about neurodegenerative diseases (Ahmed et al., 2016; Goedert et al., 2017; Guest et al., 2011; Iturria-Medina et al., 2014; Raj et al., 2012; Seeley et al., 2009; Warren et al., 2013, 2012; Zhou et al., 2012), but has also been putatively extended to psychiatric conditions. In fact, insoluble aggregates of disrupted-in-schizophrenia 1 (DISC1) were associated to sporadic cases of schizophrenia, bipolar disorder and depression (Korth, 2012; Leliveld et al., 2008). In addition, *in vitro* studies have demonstrated that aggregates of DISC1 are able to transfer between cells via tunnelling nanotubes (Zhu et al., 2017). These aggregates can selectively affect dopaminergic brain functioning at pre-synaptic and post-synaptic level (Dahoun et al., 2017; Tropea et al., 2018), and, as they are related to oxidative stress (Trossbach et al., 2016), the transneuronal spread hypothesis is not incompatible with the metabolic stress model. On the contrary, both pathological mechanisms might be necessary for a hub to be damaged (Saxena and Caroni, 2011). According to this view, brain hubs are more vulnerable to deterioration, and in turn they can spread the alterations to the connected areas. Furthermore, given their high degree of connectivity and their role of integration of different clusters, hubs are generally linked to many distant regions, and this makes them ideal for spreading pathological alterations along several long-range connections.

It has also been suggested that alterations could propagate by means of a trophic factor release failure (Fornito et al., 2015). In other words, the areas connected to the damaged region might suffer morphological GM decreases because they cease to receive trophic factors from them (Chao, 2003), or might reduce activity because of the lack of inputs, which could disrupt their activity-dependent trophic factors synthesis and release (Blöchl and Thoenen, 1995; Gall and Isackson, 1989; Kohara et al., 2001), thus leading to a cascade of anatomical decreases. A disruption of the balance of the trophic mechanism, in form of an enhanced trophic release or a lack of growth-inhibitory signals (Perlson et al., 2010), may also account for the generations of networks of GM increases. Of course, the effect of morphometric increase might be of iatrogenic nature (Hafeman et al., 2012; Navari and Dazzan, 2009); however, this general explanation is not able to take into account the similarity between hubs of long-distance increase to that of FC. Thus, we suggest that a mixture of metabolic stress, toxic spread, trophic factor release disruption and shared vulnerability to genetic and environmental factors influence the distribution of both decrease and increase co-alteration along connectivity pathways (Cauda et al., 2019). Therefore, brain hubs might be not only extremely vulnerable to disorders but also substantially responsible for the long-range spread of morphological changes.

### 4.3 Analyses of schizophrenia and Alzheimer’s disease

With regard to schizophrenia (Fig. 4), areas with long-distance co-alterations of GM decreases have auditory and linguistic roles; a finding that is in accordance with the auditory hallucinations affecting a portion of these patients (García-Martí et al., 2008; Modinos et al., 2009; Neckelmann et al., 2006; Plaze et al., 2006). Also the caudate and the MTL exhibit long-distance co-alterations, as well as the SN, which is in line with the involvement of this network in the disease (Cauda et al., 2018a; Kapur, 2003; Liddle et al., 2016; Manoliu et al., 2014; Palaniyappan et al., 2013; Palaniyappan and Liddle, 2012; Uddin, 2015; White et al., 2010). Long co-alterations of GM increases are found especially in the left putamen, which is coherent with a study that found an increased putamen characterized by leftward asymmetry in schizophrenic patients (Okada et al., 2016).

With regard to Alzheimer’s disease (Fig. 5), long-distance co-alterations of GM increases and GM decreases are found especially in the left hippocampal formation, which is consistent with the assumption that the hippocampus may be the pathological epicentre, as well as with the observation that the left hemisphere is more affected than the right one (Braak et al., 1993; Buckner, 2005; Janke et al., 2001; Loewenstein et al., 1989; Manuello et al., 2018; Pievani et al., 2011; Thompson et al., 2007, 2003, 2001). Other regions with long-distance co-alterations of GM decreases are the caudate and the insula, with a major involvement in the left hemisphere. The finding that the left MTL is involved in long-distance co-alterations both of GM increases and of GM decreases in intriguing, and might be putatively explained by the effect of compensation, as if this region could be engaged in a system of increases that try to counteract to the damages induced by the disease. Although small posteromedial cortical clusters can be appreciated in the map of GM decreases (Fig. 5), the absence of large significant neocortical clusters might be due to the modest sensitivity of VBM to changes in the cortical ribbon compared to those in the hippocampal region (Diaz-De-Grenu et al., 2014). As we have seen, the measurement of the mean physical distance of co-alterations can identify clinically relevant areas, as they are associated with typical symptoms of brain disorders, and even the epicenters of pathological diffusion. In this regard, it should be emphasized that our analysis is considerably different from other volumetric studies: our methodology does not evaluate the size and number of alterations, but only if an alteration is associated with nearby or distant modifications. The two measures, therefore, do not necessarily converge, and the fact that our findings are in line with those provided by the scientific literature is a strong indication that the mean physical distance of co-alterations is an insightful instrument for detecting pathological features of diseases.

### 4.5 Limitations

A possible limitation of our analyses is that VOIs were defined on the basis of an anatomical atlas; therefore, they may fail to account for more fine-grained distinctions in heterogeneous regions. This choice aimed to achieve a higher statistical power, as a voxel-wise technique may leave some voxels uncovered by a sufficient number of samples. It could be argued that in a parcellation the size of the ROIs determines the minimum spatial resolution for the detection of a hub, but the use of a parcellation with small volumes would have reduced the statistical power in under-represented brain regions. We therefore chose to use an atlas that previously proved itself to fit the functional connectivity better than artifactual parcellations (Mancuso et al., 2019). Still, choosing a different anatomical atlas might have produced slightly divergent results.

Another limitation concerns the practical unfeasibility to derive from the BrainMap repository data about the medication status of the large database of patients that entered this meta-analysis. Moreover, our search did not differentiate between gender and age. Therefore, analyses were unable to evaluate the effects of such variables on measuring the mean physical distance of co-alterations. Given the effect of some psychotropic drug of GM volume (Hafeman et al., 2012; Navari and Dazzan, 2009), and that age and sex have been recently reported to be associated with asymmetries in cortical thickness (Guadalupe et al., 2017; Kong et al., 2018), it could be worth investigating how the symmetry of the maps of mean physical distance of co-alterations related to GM increases and GM decreases can differ with respect to these variables.

Finally, we calculated the distance between two areas considering the length of the straight line connecting their centroids, but due to curvilinear anatomical constraints the real axonal path linking them is supposed to be probably longer. Moreover, the structural connection between two regions could be indirect, so that the topological distance too could have been useful to provide information about their proximity. However, we were mostly interested in observing whether or not areas are co-altered with far away regions (e.g. if a frontal area is co-altered only with the frontal cortex or even with regions of different lobes). From this perspective, the accurate estimation of the length of the path connecting two regions is not particularly important, though it could be valuable for the transneuronal spread hypothesis.

## 6. Conclusion

This study provides the maps of the regions characterized by long mean physical distance of co-alterations related to both GM decreases and GM increases. Our approach has produced four important findings for the understanding of brain diseases. First, when the brain is affected by a pathological process, the anterior and dorsal PFC tend to be involved in a network of long-distance co-alterations, and thus they are to be considered as key hubs of pathology. Second, areas of the left hemisphere seem to be prevalently co-altered in GM decreases, while areas of the right hemisphere appear to be more co-altered in GM increases. This suggests that the two sides of the brain are differently affected by pathological processes. Third, on the basis of the analyses of schizophrenia and Alzheimer’s disease, we have found that the measurement of the mean physical distance between co-alterations is able identify the areas that are clinically relevant for the diseases. Lastly, hubs of long-distance co-alteration are similar to those of functional connectivity DC, suggesting a relation between co-alteration and normative connectivity. The mean physical distance of co-alteration, therefore, proves itself to be a useful index capable of providing new insights into the distribution patterns of morphological alterations caused by brain disorders.

## Funding

This study was supported by Fondazione Sanpaolo, Turin (Cauda F. PI).

## Competing interest

The authors declare no competing interest.

## Notes

#### Summary of Updates

The reference section was erroneously edited by the software: 3 different series of reference were listed.

## References

Achard, S., Salvador, R., Whitcher, B., Suckling, J., Bullmore, E., 2006. A Resilient, Low-Frequency, Small-World Human Brain Functional Network with Highly Connected Association Cortical Hubs. J. Neurosci. 26, 63–72. https://doi.org/10.1523/JNEUROSCI.3874-05.2006

Ahmed, R.M., Devenney, E.M., Irish, M., Ittner, A., Naismith, S., Ittner, L.M., Rohrer, J.D., Halliday, G.M., Eisen, A., Hodges, J.R., Kiernan, M.C., 2016. Neuronal network disintegration: Common pathways linking neurodegenerative diseases. J. Neurol. Neurosurg. Psychiatry 87, 1234–1241. https://doi.org/10.1136/jnnp-2014-308350

Alexander-Bloch, A.F., Vértes, P.E., Stidd, R., Lalonde, F., Clasen, L., Rapoport, J., Giedd, J., Bullmore, E.T., Gogtay, N., 2013. The anatomical distance of functional connections predicts brain network topology in health and schizophrenia. Cereb. Cortex 23, 127–138. https://doi.org/10.1093/cercor/bhr388

Andrews-Hanna, J.R., Reidler, J.S., Sepulcre, J., Poulin, R., Buckner, R.L., 2010. Functional-Anatomic Fractionation of the Brain’s Default Network. Neuron 65, 550–562. https://doi.org/10.1016/j.neuron.2010.02.005

Bassett, D.S., Bullmore, E., Verchinski, B.A., Mattay, V.S., Weinberger, D.R., Meyer-Lindenberg, A., 2008. Hierarchical Organization of Human Cortical Networks in Health and Schizophrenia. J. Neurosci. 28, 9239–9248. https://doi.org/10.1523/JNEUROSCI.1929-08.2008

Blöchl, A., Thoenen, H., 1995. Characterization of Nerve Growth Factor (NGF) Release from Hippocampal Neurons: Evidence for a Constitutive and an Unconventional Sodium-dependent Regulated Pathway. Eur. J. Neurosci. 7, 1220–1228. https://doi.org/10.1111/j.1460-9568.1995.tb01112.x

Braak, H., Braak, E., Bohl, J., 1993. Staging of Alzheimer-Related Cortical Destruction. Eur. Neurol. 33, 403–408. https://doi.org/10.1159/000116984

Buckholtz, J.W., Meyer-Lindenberg, A., 2012. Psychopathology and the Human Connectome: Toward a Transdiagnostic Model of Risk For Mental Illness. Neuron 74, 990–1004. https://doi.org/10.1016/j.neuron.2012.06.002

Buckner, R.L., 2005. Molecular, Structural, and Functional Characterization of Alzheimer’s Disease: Evidence for a Relationship between Default Activity, Amyloid, and Memory. J. Neurosci. 25, 7709–7717. https://doi.org/10.1523/jneurosci.2177-05.2005

Buckner, R.L., Andrews-Hanna, J.R., Schacter, D.L., 2008. The brain’s default network: Anatomy, function, and relevance to disease. Ann. N. Y. Acad. Sci. 1124, 1–38. https://doi.org/10.1196/annals.1440.011

Buckner, R.L., Sepulcre, J., Talukdar, T., Krienen, F.M., Liu, H., Hedden, T., Andrews-Hanna, J.R., Sperling, R.A., Johnson, K.A., 2009. Cortical Hubs Revealed by Intrinsic Functional Connectivity: Mapping, Assessment of Stability, and Relation to Alzheimer’s Disease. J. Neurosci. 29, 1860–1873. https://doi.org/10.1523/JNEUROSCI.5062-08.2009

Bullmore, E., Sporns, O., 2012. The economy of brain network organization. Nat. Rev. Neurosci. 13, 336–349. https://doi.org/10.1038/nrn3214

Cauda, F., Costa, T., Nani, A., Fava, L., Palermo, S., Bianco, F., Duca, S., Tatu, K., Keller, R., 2017. Are schizophrenia, autistic, and obsessive spectrum disorders dissociable on the basis of neuroimaging morphological findings?: A voxel-based meta-analysis. Autism Res. 10, 1079– 1095. https://doi.org/10.1002/aur.1759

Cauda, F., Costa, T., Palermo, S., D’Agata, F., Diano, M., Bianco, F., Duca, S., Keller, R., 2014. Concordance of white matter and gray matter abnormalities in autism spectrum disorders: A voxel-based meta-analysis study. Hum. Brain Mapp. 35, 2073–2098. https://doi.org/10.1002/hbm.22313

Cauda, F., Mancuso, L., Nani, A., Costa, T., 2019. Heterogeneous neuroimaging findings, damage propagation and connectivity: an integrative view. Brain 142, 1–3. https://doi.org/10.1093/brain/awz080

Cauda, F., Nani, A., Costa, T., Palermo, S., Tatu, K., Manuello, J., Duca, S., Fox, P.T., Keller, R., 2018a. The morphometric co-atrophy networking of schizophrenia, autistic and obsessive spectrum disorders. Hum. Brain Mapp. 39, 1898–1928. https://doi.org/10.1002/hbm.23952

Cauda, F., Nani, A., Manuello, J., Premi, E., Palermo, S., Tatu, K., Duca, S., Fox, P.T., Costa, T., 2018b. Brain structural alterations are distributed following functional, anatomic and genetic connectivity. Brain 141, 3211–3232. https://doi.org/10.1093/brain/awy252

Chao, M. V., 2003. Neurotrophins and their receptors: A convergence point for many signalling pathways. Nat. Rev. Neurosci. 4, 299–309. https://doi.org/10.1038/nrn1078

Crossley, N.A., Mechelli, A., Scott, J., Carletti, F., Fox, P.T., Mcguire, P., Bullmore, E.T., 2014. The hubs of the human connectome are generally implicated in the anatomy of brain disorders. Brain 137, 2382–2395. https://doi.org/10.1093/brain/awu132

Dahoun, T., Trossbach, S. V., Brandon, N.J., Korth, C., Howes, O.D., 2017. The impact of Disrupted-in-Schizophrenia 1 (DISC1) on the dopaminergic system: A systematic review. Transl. Psychiatry 7, e1015–15. https://doi.org/10.1038/tp.2016.282

de Haan, W., Mott, K., van Straaten, E.C.W., Scheltens, P., Stam, C.J., 2012. Activity Dependent Degeneration Explains Hub Vulnerability in Alzheimer’s Disease. PLoS Comput. Biol. 8. https://doi.org/10.1371/journal.pcbi.1002582

Diaz-De-Grenu, L.Z., Acosta-Cabronero, J., Chong, Y.F.V., Pereira, J.M.S., Sajjadi, S.A., Williams, G.B., Nestor, P.J., 2014. A brief history of voxel-based grey matter analysis in Alzheimer’s disease. J. Alzheimer’s Dis. 38, 647–659. https://doi.org/10.3233/JAD-130362

Eickhoff, S., Laird, A., Grefkes, C., Wang, L.E., Zilles, K., Fox, P.T., 2009. Coordinate-based ALE meta-analysis of neuroimaging data: a random-effects approach based on empirical estimates of spatial uncertainty. Hum. Brain Mapp. 30, 2907–2926. https://doi.org/10.1002/hbm.20718.Coordinate-based

Eickhoff, S.B., Bzdok, D., Laird, A.R., Kurth, F., Fox, P.T., 2012. Activation likelihood estimation revisited. Neuroimage 59, 2349–2361. https://doi.org/10.1016/j.neuroimage.2011.09.017.Activation

Esfahlani, F.Z., Bertolero, M., Bassett, D., Betzel, R., 2019. Space-independent community and hub structure of functional brain networks. bioRxiv 590935. https://doi.org/10.1101/590935

Evans, A.C., 2013. Networks of anatomical covariance. Neuroimage 80, 489–504. https://doi.org/10.1016/j.neuroimage.2013.05.054

Fornito, A., Zalesky, A., Breakspear, M., 2015. The connectomics of brain disorders. Nat. Rev. Neurosci. 16, 159–172. https://doi.org/10.1038/nrn3901

Fox, P.T., Laird, A.R., Fox, S.P., Fox, P.M., Uecker, A.M., Crank, M., Koenig, S.F., Lancaster, J.L., 2005. BrainMap taxonomy of experimental design: Description and evaluation. Hum. Brain Mapp. 25, 185–198. https://doi.org/10.1002/hbm.20141

Fox, P.T., Lancaster, J.L., 2002. Mapping context and content: The BrainMap model. Nat. Rev. Neurosci. 3, 319–321. https://doi.org/10.1038/nrn789

Gall, C.M., Isackson, P.J., 1989. Limbic seizures increase neuronal production of messenger RNA for nerve growth factor. Science (80-.). https://doi.org/10.1126/science.2549634

García-Martí, G., Aguilar, E.J., Lull, J.J., Martí-Bonmatí, L., Escartí, M.J., Manjón, J. V., Moratal, D., Robles, M., Sanjuán, J., 2008. Schizophrenia with auditory hallucinations: A voxel-based morphometry study. Prog. Neuro-Psychopharmacology Biol. Psychiatry 32, 72–80. https://doi.org/10.1016/j.pnpbp.2007.07.014

Goedert, M., Masuda-Suzukake, M., Falcon, B., 2017. Like prions: The propagation of aggregated tau and α-synuclein in neurodegeneration. Brain 140, 266–278. https://doi.org/10.1093/brain/aww230

Goodkind, M., Eickhoff, S.B., Oathes, D.J., Jiang, Y., Chang, A., Jones-Hagata, L.B., Ortega, B.N., Zaiko, Y. V., Roach, E.L., Korgaonkar, M.S., Grieve, S.M., Galatzer-Levy, I., Fox, P.T., Etkin, A., 2015. Identification of a common neurobiological substrate for mental Illness. JAMA Psychiatry 72, 305–315. https://doi.org/10.1001/jamapsychiatry.2014.2206

Green, S., Higgins, J.P., Alderson, P., Clarke, M., Mulrow, C.D., Oxman, A.D., 2008. Introduction, in: Higgins, J.P., Green, S. (Eds.), Cochrane Handbook for Systematic Reviews of Interventions. John Wiley & Sons, Ltd, Chichester, UK, pp. 1–9. https://doi.org/10.1002/9780470712184.ch1

Guadalupe, T., Mathias, S.R., vanErp, T.G.M., Whelan, C.D., Zwiers, M.P., Abe, Y., Abramovic, L., Agartz, I., Andreassen, O.A., Arias-Vásquez, A., Aribisala, B.S., Armstrong, N.J., Arolt, V., Artiges, E., Ayesa-Arriola, R., Baboyan, V.G., Banaschewski, T., Barker, G., Bastin, M.E., Baune, B.T., Blangero, J., Bokde, A.L.W., Boedhoe, P.S.W., Bose, A., Brem, S., Brodaty, H., Bromberg, U., Brooks, S., Büchel, C., Buitelaar, J., Calhoun, V.D., Cannon, D.M., Cattrell, A., Cheng, Y., Conrod, P.J., Conzelmann, A., Corvin, A., Crespo-Facorro, B., Crivello, F., Dannlowski, U., de Zubicaray, G.I., de Zwarte, S.M.C., Deary, I.J., Desrivières, S., Doan, N.T., Donohoe, G., Dørum, E.S., Ehrlich, S., Espeseth, T., Fernández, G., Flor, H., Fouche, J.P., Frouin, V., Fukunaga, M., Gallinat, J., Garavan, H., Gill, M., Suarez, A.G., Gowland, P., Grabe, H.J., Grotegerd, D., Gruber, O., Hagenaars, S., Hashimoto, R., Hauser, T.U., Heinz, A., Hibar, D.P., Hoekstra, P.J., Hoogman, M., Howells, F.M., Hu, H., Hulshoff Pol, H.E., Huyser, C., Ittermann, B., Jahanshad, N., Jönsson, E.G., Jurk, S., Kahn, R.S., Kelly, S., Kraemer, B., Kugel, H., Kwon, J.S., Lemaitre, H., Lesch, K.P., Lochner, C., Luciano, M., Marquand, A.F., Martin, N.G., Martínez-Zalacaín, I., Martinot, J.L., Mataix-Cols, D., Mather, K., McDonald, C., McMahon, K.L., Medland, S.E., Menchón, J.M., Morris, D.W., Mothersill, O., Maniega, S.M., Mwangi, B., Nakamae, T., Nakao, T., Narayanaswaamy, J.C., Nees, F., Nordvik, J.E., Onnink, A.M.H., Opel, N., Ophoff, R., Paillère Martinot, M.L., Papadopoulos Orfanos, D., Pauli, P., Paus, T., Poustka, L., Reddy, J.Y., Renteria, M.E., Roiz-Santiáñez, R., Roos, A., Royle, N.A., Sachdev, P., Sánchez-Juan, P., Schmaal, L., Schumann, G., Shumskaya, E., Smolka, M.N., Soares, J.C., Soriano-Mas, C., Stein, D.J., Strike, L.T., Toro, R., Turner, J.A., Tzourio-Mazoyer, N., Uhlmann, A., Hernández, M.V., van den Heuvel, O.A., van der Meer, D., van Haren, N.E.M., Veltman, D.J., Venkatasubramanian, G., Vetter, N.C., Vuletic, D., Walitza, S., Walter, H., Walton, E., Wang, Z., Wardlaw, J., Wen, W., Westlye, L.T., Whelan, R., Wittfeld, K., Wolfers, T., Wright, M.J., Xu, J., Xu, X., Yun, J.Y., Zhao, J.J., Franke, B., Thompson, P.M., Glahn, D.C., Mazoyer, B., Fisher, S.E., Francks, C., 2017. Human subcortical brain asymmetries in 15,847 people worldwide reveal effects of age and sex. Brain Imaging Behav. 11, 1497–1514. https://doi.org/10.1007/s11682-016-9629-z

Guest, W.C., Silverman, J.M., Pokrishevsky, E., O’Neill, M.A., Grad, L.I., Cashman, N.R., 2011. Generalization of the Prion Hypothesis to Other Neurodegenerative Diseases: An Imperfect Fit. J. Toxicol. Environ. Heal. Part A 74, 1433–1459. https://doi.org/10.1080/15287394.2011.618967

Hafeman, D.M., Chang, K.D., Garrett, A.S., Sanders, E.M., Phillips, M.L., 2012. Effects of medication on neuroimaging findings in bipolar disorder: An updated review. Bipolar Disord. 14, 375–410. https://doi.org/10.1111/j.1399-5618.2012.01023.x

He, Y., Chen, Z.J., Evans, A.C., 2007. Small-world anatomical networks in the human brain revealed by cortical thickness from MRI. Cereb. Cortex 17, 2407–2419. https://doi.org/10.1093/cercor/bhl149

Iturria-Medina, Y., Sotero, R.C., Toussaint, P.J., Evans, A.C., 2014. Epidemic Spreading Model to Characterize Misfolded Proteins Propagation in Aging and Associated Neurodegenerative Disorders. PLoS Comput. Biol. 10. https://doi.org/10.1371/journal.pcbi.1003956

Janke, A.L., De Zubicaray, G., Rose, S.E., Griffin, M., Chalk, J.B., Galloway, G.J., 2001. 4D deformation modeling of cortical disease progression in Alzheimer’s dementia. Magn. Reson. Med. 46, 661–666. https://doi.org/10.1002/mrm.1243

Kapur, S., 2003. Psychosis as a state of aberrant salience: A framework linking biology, phenomenology, and pharmacology in schizophrenia. Am. J. Psychiatry 160, 13–23. https://doi.org/10.1176/appi.ajp.160.1.13

Kohara, K., Kitamura, A., Morishima, M., Tsumoto, T., 2001. Activity-dependent transfer of brain-derived neurotrophic factor to postsynaptic neurons. Science (80-.). 291, 2419–2423. https://doi.org/10.1126/science.1057415

Kong, X.-Z., Mathias, S.R., Guadalupe, T., Glahn, D.C., Franke, B., Crivello, F., Tzourio-Mazoyer, N., Fisher, S.E., Thompson, P.M., Francks, C., 2018. Mapping cortical brain asymmetry in 17,141 healthy individuals worldwide via the ENIGMA Consortium. Proc. Natl. Acad. Sci. 115, E5154–E5163. https://doi.org/10.1073/pnas.1718418115

Korth, C., 2012. Aggregated proteins in schizophrenia and other chronic mental diseases. Prion 6, 134–141. https://doi.org/10.4161/pri.18989

Laird, A.R., Lancaster, J.L., Fox, P.T., 2005. BrainMap: The Social Evolution of a Human Brain Mapping Database. Neuroinformatics 3, 065–078. https://doi.org/10.1385/NI:3:1:065

Lancaster, J.L., Tordesillas-Gutiérrez, D., Martinez, M., Salinas, F., Evans, A., Zilles, K., Mazziotta, J.C., Fox, P.T., 2007. Bias between MNI and Talairach coordinates analyzed using the ICBM-152 brain template. Hum. Brain Mapp. 28, 1194–1205. https://doi.org/10.1002/hbm.20345

Lancaster, J.L., Woldorff, M.G., Parsons, L.M., Liotti, M., Freitas, C.S., Rainey, L., Kochunov, P. V., Nickerson, D., Mikiten, S.A., Fox, P.T., 2000. Automated Talairach Atlas labels for functional brain mapping. Hum. Brain Mapp. 10, 120–131. https://doi.org/10.1002/1097-0193(200007)10:3<120::AID-HBM30>3.0.CO;2-8

Laughlin, S.B., Sejnowski, T.J., 2003. Communication in neuronal networks. Science (80-.). 301, 1870–1874. https://doi.org/10.1126/science.1089662

Leliveld, S.R., Bader, V., Hendriks, P., Prikulis, I., Sajnani, G., Requena, J.R., Korth, C., 2008. Insolubility of Disrupted-in-Schizophrenia 1 Disrupts Oligomer-Dependent Interactions with Nuclear Distribution Element 1 and Is Associated with Sporadic Mental Disease. J. Neurosci. 28, 3839–3845. https://doi.org/10.1523/JNEUROSCI.5389-07.2008

Liao, X.H., Xia, M.R., Xu, T., Dai, Z.J., Cao, X.Y., Niu, H.J., Zuo, X.N., Zang, Y.F., He, Y., 2013. Functional brain hubs and their test-retest reliability: A multiband resting-state functional MRI study. Neuroimage 83, 969–982. https://doi.org/10.1016/j.neuroimage.2013.07.058

Liberati, A., Altman, D.G., Tetzlaff, J., Mulrow, C., Gotzsche, P.C., Ioannidis, J.P.A., Clarke, M., Devereaux, P.J., Kleijnen, J., Moher, D., 2009. The PRISMA statement for reporting systematic reviews and meta-analyses of studies that evaluate healthcare interventions: explanation and elaboration. BMJ 339, b2700–b2700. https://doi.org/10.1136/bmj.b2700

Liddle, E.B., Price, D., Palaniyappan, L., Brookes, M.J., Robson, S.E., Hall, E.L., Morris, P.G., Liddle, P.F., 2016. Abnormal salience signaling in schizophrenia: The role of integrative beta oscillations. Hum. Brain Mapp. 37, 1361–1374. https://doi.org/10.1002/hbm.23107

Loewenstein, D.A., Barker, W.W., Chang, J.Y., Apicella, A., Yoshii, F., Kothari, P., Levin, B., Duara, R., 1989. Predominant left hemisphere metabolic dysfunction in dementia. Arch. Neurol. 46, 146–152. https://doi.org/10.1001/archneur.1989.00520380046012

Mancuso, L., Costa, T., Nani, A., Manuello, J., Liloia, D., Gelmini, G., Panero, M., Duca, S., Cauda, F., 2019. The homotopic connectivity of the functional brain: a meta-analytic approach. Sci. Rep. 9, 3346. https://doi.org/10.1038/s41598-019-40188-3

Mandelli, M.L., Vilaplana, E., Brown, J.A., Hubbard, H.I., Binney, R.J., Attygalle, S., Santos-Santos, M.A., Miller, Z.A., Pakvasa, M., Henry, M.L., Rosen, H.J., Henry, R.G., Rabinovici, G.D., Miller, B.L., Seeley, W.W., Gorno-Tempini, M.L., 2016. Healthy brain connectivity predicts atrophy progression in non-fluent variant of primary progressive aphasia. Brain 139, 2778–2791. https://doi.org/10.1093/brain/aww195

Manoliu, A., Riedl, V., Zherdin, A., Mühlau, M., Schwerthöffer, D., Scherr, M., Peters, H., Zimmer, C., Förstl, H., Bäuml, J., Wohlschläger, A.M., Sorg, C., 2014. Aberrant dependence of default mode/central executive network interactions on anterior insular salience network activity in schizophrenia. Schizophr. Bull. 40, 428–437. https://doi.org/10.1093/schbul/sbt037

Manuello, J., Nani, A., Premi, E., Borroni, B., Costa, T., Tatu, K., Liloia, D., Duca, S., Cauda, F., 2018. The pathoconnectivity profile of Alzheimer’s disease: A morphometric coalteration network analysis. Front. Neurol. 8, 1–15. https://doi.org/10.3389/fneur.2017.00739

McTeague, L.M., Goodkind, M.S., Etkin, A., 2016. Transdiagnostic impairment of cognitive control in mental illness. J. Psychiatr. Res. 83, 37–46. https://doi.org/10.1016/j.jpsychires.2016.08.001

Modinos, G., Vercammen, A., Mechelli, A., Knegtering, H., McGuire, P.K., Aleman, A., 2009. Structural covariance in the hallucinating brain: A voxel-based morphometry study. J. Psychiatry Neurosci. 34, 465–469. https://doi.org/10.1016/S1701-2163(16)32305-2

Moher, D., Liberati, A., Tetzlaff, J., Altman, D.G., 2009. Preferred Reporting Items for Systematic Reviews and Meta-Analyses: The PRISMA Statement. J. Clin. Epidemiol. 62, 1006–1012. https://doi.org/10.1016/j.jclinepi.2009.06.005

Navari, S., Dazzan, P., 2009. Do antipsychotic drugs affect brain structure? A systematic and critical review of MRI findings. Psychol. Med. 39, 1763–1777. https://doi.org/10.1017/S0033291709005315

Neckelmann, G., Specht, K., Lund, A., Ersland, L., Smievoll, A.I., Neckelmann, D., Hugdahl, K., 2006. MR morphometry analysis of grey matter volume reduction in schizophrenia: Association with hallucinations. Int. J. Neurosci. 116, 9–23. https://doi.org/10.1080/00207450690962244

Okada, N., Fukunaga, M., Yamashita, F., Koshiyama, D., Yamamori, H., Ohi, K., Yasuda, Y., Fujimoto, M., Watanabe, Y., Yahata, N., Nemoto, K., Hibar, D.P., Van Erp, T.G.M., Fujino, H., Isobe, M., Isomura, S., Natsubori, T., Narita, H., Hashimoto, N., Miyata, J., Koike, S., Takahashi, T., Yamasue, H., Matsuo, K., Onitsuka, T., Iidaka, T., Kawasaki, Y., Yoshimura, R., Watanabe, Y., Suzuki, M., Turner, J.A., Takeda, M., Thompson, P.M., Ozaki, N., Kasai, K., Hashimoto, R., 2016. Abnormal asymmetries in subcortical brain volume in schizophrenia. Mol. Psychiatry 21, 1460–1466. https://doi.org/10.1038/mp.2015.209

Palaniyappan, L., Liddle, P.F., 2012. Does the salience network play a cardinal role in psychosis? An emerging hypothesis of insular dysfunction. J. Psychiatry Neurosci. 37, 17–27. https://doi.org/10.1503/jpn.100176

Palaniyappan, L., White, T.P., Liddle, P.F., 2013. The Concept of Salience Network Dysfunction in Schizophrenia: From Neuroimaging Observations to Therapeutic Opportunities. Curr. Top. Med. Chem. 12, 2324–2338. https://doi.org/10.2174/1568026611212210005

Patel, R.S., Bowman, F.D., Rilling, J.K., 2006. A Bayesian approach to determining connectivity of the human brain. Hum. Brain Mapp. 27, 267–276. https://doi.org/10.1002/hbm.20182

Perlson, E., Maday, S., Fu, M. meng, Moughamian, A.J., Holzbaur, E.L.F., 2010. Retrograde axonal transport: Pathways to cell death? Trends Neurosci. 33, 335–344. https://doi.org/10.1016/j.tins.2010.03.006

Pievani, M., de Haan, W., Wu, T., Seeley, W.W., Frisoni, G.B., 2011. Functional network disruption in the degenerative dementias. Lancet Neurol. 10, 829–843. https://doi.org/10.1016/S1474-4422(11)70158-2

Plaze, M., Bartrés-Faz, D., Martinot, J.L., Januel, D., Bellivier, F., De Beaurepaire, R., Chanraud, S., Andoh, J., Lefaucheur, J.P., Artiges, E., Pallier, C., Paillère-Martinot, M.L., 2006. Left superior temporal gyrus activation during sentence perception negatively correlates with auditory hallucination severity in schizophrenia patients. Schizophr. Res. 87, 109–115. https://doi.org/10.1016/j.schres.2006.05.005

Raj, A., Kuceyeski, A., Weiner, M., 2012. A Network Diffusion Model of Disease Progression in Dementia. Neuron 73, 1204–1215. https://doi.org/10.1016/j.neuron.2011.12.040

Rubinov, M., Sporns, O., 2010. Complex network measures of brain connectivity: Uses and interpretations. Neuroimage 52, 1059–1069. https://doi.org/10.1016/j.neuroimage.2009.10.003

Saxena, S., Caroni, P., 2011. Selective Neuronal Vulnerability in Neurodegenerative Diseases: From Stressor Thresholds to Degeneration. Neuron 71, 35–48. https://doi.org/10.1016/j.neuron.2011.06.031

Scarpazza, C., Tognin, S., Frisciata, S., Sartori, G., Mechelli, A., 2015. False positive rates in Voxel-based Morphometry studies of the human brain: Should we be worried? Neurosci. Biobehav. Rev. 52, 49–55. https://doi.org/10.1016/j.neubiorev.2015.02.008

Seeley, W.W., Crawford, R.K., Zhou, J., Miller, B.L., Greicius, M.D., 2009. Neurodegenerative Diseases Target Large-Scale Human Brain Networks. Neuron 62, 42–52. https://doi.org/10.1016/j.neuron.2009.03.024

Sepulcre, J., Liu, H., Talukdar, T., Martincorena, I., Thomas Yeo, B.T., Buckner, R.L., 2010. The organization of local and distant functional connectivity in the human brain. PLoS Comput. Biol. 6, 1–15. https://doi.org/10.1371/journal.pcbi.1000808

Sheline, Y.I., Barch, D.M., Price, J.L., Rundle, M.M., Vaishnavi, S.N., Snyder, A.Z., Mintun, M.A., Wang, S., Coalson, R.S., Raichle, M.E., 2009. The default mode network and self-referential processes in depression. Proc. Natl. Acad. Sci. 106, 1942–1947. https://doi.org/10.1073/pnas.0812686106

Sporns, O., Zwi, J.D., 2004. The Small World of the Cerebral Cortex. Neuroinformatics 2, 145– 162. https://doi.org/10.1385/NI:2:2:145

Tatu, K., Costa, T., Nani, A., Diano, M., Quarta, D.G., Duca, S., Apkarian, A.V., Fox, P.T., Cauda, F., 2018. How do morphological alterations caused by chronic pain distribute across the brain? A meta-analytic co-alteration study. NeuroImage Clin. 18, 15–30. https://doi.org/10.1016/j.nicl.2017.12.029

Thompson, P.M., Hayashi, K.M., de Zubicaray, G., Janke, A.L., Rose, S.E., Semple, J., Herman, D., Hong, M.S., Dittmer, S.S., Doddrell, D.M., Toga, A.W., 2003. Dynamics of Gray Matter Loss in Alzheimer’s Disease. J. Neurosci. 23, 994–1005. https://doi.org/10.1523/JNEUROSCI.23-03-00994.2003

Thompson, P.M., Hayashi, K.M., Dutton, R.A., Chiang, M.-C., Leow, A.D., SowellL, E.R., de Zubicaray, G., Becker, J.T., Lopez, O.L., Aizenstein, H.J., Toga, A.W., 2007. Tracking Alzheimer’s Disease. Ann. N. Y. Acad. Sci. 1097, 183–214. https://doi.org/10.1196/annals.1379.017

Thompson, P.M., Mega, M.S., Woods, R.P., Zoumalan, C.I., Lindshield, C.J., Blanton, R.E., Moussai, J., Holmes, C.J., Cummings, J.L., Toga, A.W., 2001. Cortical Change in Alzheimer’s Disease Detected with a Disease-specific Population-based Brain Atlas. Cereb. Cortex 11, 1– 16. https://doi.org/10.1093/cercor/11.1.1

Tropea, D., Hardingham, N., Millar, K., Fox, K., 2018. Mechanisms underlying the role of DISC1 in synaptic plasticity. J. Physiol. 596, 2747–2771. https://doi.org/10.1113/JP274330

Trossbach, S. V., Bader, V., Hecher, L., Pum, M.E., Masoud, S.T., Prikulis, I., Schäble, S., De Souza Silva, M.A., Su, P., Boulat, B., Chwiesko, C., Poschmann, G., Stühler, K., Lohr, K.M., Stout, K.A., Oskamp, A., Godsave, S.F., Müller-Schiffmann, A., Bilzer, T., Steiner, H., Peters, P.J., Bauer, A., Sauvage, M., Ramsey, A.J., Miller, G.W., Liu, F., Seeman, P., Brandon, N.J., Huston, J.P., Korth, C., 2016. Misassembly of full-length Disrupted-in-Schizophrenia 1 protein is linked to altered dopamine homeostasis and behavioral deficits. Mol. Psychiatry 21, 1561– 1572. https://doi.org/10.1038/mp.2015.194

Turkeltaub, P.E., Eickhoff, S.B., Laird, A.R., Fox, M., Wiener, M., Fox, P., 2012. Minimizing within-experiment and within-group effects in activation likelihood estimation meta-analyses. Hum. Brain Mapp. 33, 1–13. https://doi.org/10.1002/hbm.21186

Uddin, L.Q., 2015. Salience processing and insular cortical function and dysfunction, in: Nature Reviews Neuroscience. Nature Publishing Group, pp. 55–61. https://doi.org/10.1038/nrn3857

Vanasse, T.J., Fox, P.M., Barron, D.S., Robertson, M., Eickhoff, S.B., Lancaster, J.L., Fox, P.T., 2018. BrainMap VBM: An environment for structural meta-analysis. Hum. Brain Mapp. 39, 3308–3325. https://doi.org/10.1002/hbm.24078

Warren, J.D., Rohrer, J.D., Hardy, J., 2012. Disintegrating Brain Networks: From Syndromes to Molecular Nexopathies. Neuron 73, 1060–1062. https://doi.org/10.1016/j.neuron.2012.03.006

Warren, J.D., Rohrer, J.D., Schott, J.M., Fox, N.C., Hardy, J., Rossor, M.N., 2013. Molecular nexopathies: a new paradigm of neurodegenerative disease. Trends Neurosci. 36, 561–569. https://doi.org/10.1016/j.tins.2013.06.007

Watts, D.J., Strogatz, S.H., Tseng, Y.-M., 1998. Collective dynamics of ‘small-world’ networks. Nature 153, 810–817. https://doi.org/10.1038/30918

Wheeler, A.L., Felsky, D., Viviano, J.D., Stojanovski, S., Ameis, S.H., Szatmari, P., Lerch, J.P., Chakravarty, M.M., Voineskos, A.N., 2017. BDNF-Dependent Effects on Amygdala–Cortical Circuitry and Depression Risk in Children and Youth. Cereb. Cortex 28, 1760–1770. https://doi.org/10.1093/cercor/bhx086

Wheeler, A.L., Wessa, M., Szeszko, P.R., Foussias, G., Chakravarty, M.M., Lerch, J.P., DeRosse, P., Remington, G., Mulsant, B.H., Linke, J., Malhotra, A.K., Voineskos, A.N., 2015. Further neuroimaging evidence for the deficit subtype of schizophrenia: A cortical connectomics analysis. JAMA Psychiatry 72, 446–455. https://doi.org/10.1001/jamapsychiatry.2014.3020

White, T.P., Joseph, V., Francis, S.T., Liddle, P.F., 2010. Aberrant salience network (bilateral insula and anterior cingulate cortex) connectivity during information processing in schizophrenia. Schizophr. Res. 123, 105–115. https://doi.org/10.1016/j.schres.2010.07.020

World Health Organisation, 1992. The ICD-10 classification of mental and behavioural disorders: clinical descriptions and diagnostic guidelines. World Heal. Organ. https://doi.org/10.1002/1520-6505(2000)9:5<201::AID-EVAN2>3.3.CO;2-P

Zhou, J., Gennatas, E.D., Kramer, J.H., Miller, B.L., Seeley, W.W., 2012. Predicting Regional Neurodegeneration from the Healthy Brain Functional Connectome. Neuron 73, 1216–1227. https://doi.org/10.1016/j.neuron.2012.03.004

Zhu, S., Abounit, S., Korth, C., Zurzolo, C., 2017. Transfer of disrupted-in-schizophrenia 1 aggregates between neuronal-like cells occurs in tunnelling nanotubes and is promoted by dopamine. Open Biol. 7. https://doi.org/10.1098/rsob.160328

