## Supplementary Materials for "Hubs of long-distance co-alteration in brain pathology"

centrality in coalteration brain networks

A TRANSDIAGNOSTIC AND SINGLE DISEASE APPROACH

Supplementary Material

**Figure S1. PRISMA flow chart**. Overview of the selection strategy (voxel-based morphometry and functional datasets).

***Identification***

BrainMap **Structural** sector

BrainMap **Functional** sector

**VBM Query 1**

Controls > Patients

**VBM Query 2**
Controls < Patients

**Functional Query**
Healty subjects

***Screening***

**Functional Query**
13148Experiments

**VBM Query 1**

912 Experiments

**VBM Query 2**

350 Experiments

***Eligibility***

**912** Experiments assessed

for eligibility

**350** Experiments assessed

for eligibility

**13148** Experiments assessed

for eligibility

Experiments excluded:

**170** non-relevant disorders

**65** less than 8 subjects

**35** not pathology vs healthy control

Experiments excluded:

**88** non-relevant disorders

**24** less than 8 subjects

**34** not pathology vs healthy control

Experiments included if report:

a) whole-brain analysis;

b) healty sample;

c) specified paradigm of task;

d) foci of activation in TAL or MNI.

***Included***

**432 Articles**

**642 Experiments included**

**(meta-analysis)**

**135 Articles**

**204 Experiments included**

**(meta-analysis)**

**2376 Articles**

**13148 Experiments included**

**(meta-analysis)**

Supplementary Tables

**Table S1.** Distribution of the voxel-based morphometry experimental dataset of gray matter (GM) decreases with respective diagnostic labeling (ICD-10 code) (VBM Query 1).

Number of experiments (GM decreases), percentage of the total of the experiments and subjects for each selected ICD-10 class was reported.

| **ICD-10 Code** | **Exp (n)** | **Exp (%)** | **Subj (n)** |
| --- | --- | --- | --- |
| F20: Schizophrenia | 115 | 17.9 | 3807 |
| F32-F33: Major/single episode/recurrent Depressive Disorder | 63 | 9.8 | 1919 |
| G40: Epilepsy and Recurrent Seizures | 56 | 8.7 | 1426 |
| G30: Alzheimer's Disease | 53 | 8.3 | 1194 |
| G31: Other Degenerative Diseases of Nervous System | 52 | 8.1 | 838 |
| F31: Bipolar Disorder | 45 | 7.0 | 1191 |
| G35: Multiple Sclerosis | 45 | 7.0 | 1422 |
| F84: Pervasive Developmental Disorders | 29 | 4.5 | 696 |
| G20: Parkinson's Disease | 28 | 4.4 | 501 |
| F43: Reaction to Severe Stress and Adjustment Disorders | 24 | 3.7 | 367 |
| G12: Spinal Muscular Atrophy and Related Syndromes | 14 | 2.2 | 211 |
| F41: Other Anxiety Disorders | 13 | 2.0 | 306 |
| G23: Other degenerative diseases of basal ganglia | 13 | 2.0 | 212 |
| G90: Disorders Autonomic Nervous System | 13 | 2.0 | 181 |
| F42: Obsessive Compulsive Disorder | 12 | 1.9 | 377 |
| F50: Eating Disorders | 12 | 1.9 | 148 |
| G10: Huntington's Disease | 12 | 1.9 | 258 |
| F60: Specific Personality Disorders | 10 | 1.6 | 222 |
| F90: Attention Deficit/Hyperactivity Disorder | 9 | 1.4 | 139 |
| G25: Other extrapyramidal and movement disorders | 7 | 1.1 | 135 |
| G24: Dystonia | 6 | 0.9 | 89 |
| F28: Other psychotic disorder not due to a substance or known physiological condition | 3 | 0.5 | 68 |
| G93: Other disorders of brain | 3 | 0.5 | 40 |
| F80: Specific Developmental Disorders of Speech and Language | 2 | 0.3 | 22 |
| F95: Tic Disorder | 2 | 0.3 | 33 |
| F65: Paraphilias | 1 | 0.2 | 18 |
| **Total** | **642** | **100%** | **15820** |

**Table S2.** Distribution of the voxel-based morphometry experimental dataset of gray matter (GM) increases with respective diagnostic labeling (ICD-10 code) (VBM Query 2).

Number of experiments (GM increases), percentage of the total of the experiments and subjects for each selected ICD-10 class was reported.

| **ICD-10 Code** | **Exp (n)** | **Exp (%)** | **Subj (n)** |
| --- | --- | --- | --- |
| F20: Schizophrenia | 33 | 16.2 | 1175 |
| G40: Epilepsy and Recurrent Seizures | 26 | 12.7 | 713 |
| F84: Pervasive Developmental Disorders | 23 | 11.3 | 515 |
| F31: Bipolar Disorder | 20 | 9.8 | 545 |
| F32-F33: Major depressive disorder, single episode/recurrent | 19 | 9.3 | 453 |
| F42: Obsessive Compulsive Disorder | 10 | 4.9 | 287 |
| G31: Other Degenerative Diseases of Nervous System | 10 | 4.9 | 123 |
| G35: Multiple Sclerosis | 10 | 4.9 | 253 |
| G20: Parkinson's Disease | 9 | 4.4 | 139 |
| G24: Dystonia | 8 | 3.9 | 133 |
| G25: Other extrapyramidal and movement disorders | 7 | 3.4 | 125 |
| G30: Alzheimer's Disease | 7 | 3.4 | 114 |
| F41: Other Anxiety Disorders | 4 | 2.0 | 51 |
| F50: Eating Disorders | 4 | 2.0 | 54 |
| F28: Other psychotic disorder not due to a substance or known physiological condition | 3 | 1.5 | 75 |
| F80: Specific Developmental Disorders of Speech and Language | 3 | 1.5 | 48 |
| F43: Reaction to Severe Stress and Adjustment Disorders | 2 | 1.0 | 48 |
| F90: Attention Deficit/Hyperactivity Disorder | 2 | 1.0 | 27 |
| F95: Tic Disorder | 2 | 1.0 | 45 |
| G10: Huntington's Disease | 1 | 0.5 | 21 |
| G12: Spinal Muscular Atrophy and Related Syndromes | 1 | 0.5 | 22 |
| **Total** | **204** | **100%** | **4966** |

**Table S3.** Selected articles for the meta-analysis of gray matter decreases (VBM Query 1).

| **ID** | **First Author** | **Year** | **Medline** | **ICD-10 Code** |
| --- | --- | --- | --- | --- |
| 1 | Abe O | 2010 | - | F32-F33: Major depressive disorder. single episode/recurrent |
| 2 | Abell F | 1999 | 10501551 | F84: Pervasive Developmental Disorders |
| 3 | Adleman N E | 2012 | 3472043 | F31: Bipolar Disorder |
| 4 | Adler C M | 2005 | 15922309 | F31: Bipolar Disorder |
| 5 | Agosta F | 2007 | 17370339 | G12: Spinal Muscular Atrophy and Related Syndromes |
| 6 | Agosta F | 2010 | 20597976 | G23: Other degenerative diseases of basal ganglia |
| 7 | Agosta F | 2011 | 21177393 | G30: Alzheimer's Disease |
| 8 | Agosta F | 2011 | 21177393 | G31: Other Degenerative Diseases of Nervous System |
| 9 | Ahmed F | 2012 | 22948482 | F43: Reaction to Severe Stress and Adjustment Disorders |
| 10 | Ahrendts J | 2011 | 20879808 | F90: Attention Deficit/Hyperactivity Disorder |
| 11 | Alemany S | 2013 | - | F32-F33: Major depressive disorder. single episode/recurrent |
| 12 | Almeida J R C | 2009 | 19101126 | F31: Bipolar Disorder |
| 13 | Alonso_Lana S | 2016 | 4957815 | F31: Bipolar Disorder |
| 14 | Ambrosi E | 2013 | - | F31: Bipolar Disorder |
| 15 | Ananth H | 2002 | 12202269 | F20: Schizophrenia |
| 16 | Antonova E | 2005 | 16039619 | F20: Schizophrenia |
| 17 | Arnone D | 2009 | - | F32-F33: Major depressive disorder. single episode/recurrent |
| 18 | Arnone D | 2013 | 23128153 | F32-F33: Major depressive disorder. single episode/recurrent |
| 19 | Asami T | 2009 | 19560907 | F41: Other Anxiety Disorders |
| 20 | Ash S | 2011 | 21689852 | G31: Other Degenerative Diseases of Nervous System |
| 21 | Audoin B | 2004 | 15503338 | G35: Multiple Sclerosis |
| 22 | Audoin B | 2006 | 17093899 | G35: Multiple Sclerosis |
| 23 | Audoin B | 2010 | 20392976 | G35: Multiple Sclerosis |
| 24 | Audoin B | 2007 | 17463071 | G35: Multiple Sclerosis |
| 25 | Barbeau E | 2008 | 18191160 | G31: Other Degenerative Diseases of Nervous System |
| 26 | Baron J C | 2001 | 11467904 | G30: Alzheimer's Disease |
| 27 | Bassitt D P | 2007 | 16960651 | F20: Schizophrenia |
| 28 | Baxter L C | 2006 | 16914835 | G30: Alzheimer's Disease |
| 29 | Bell-McGinty S | 2005 | 16157746 | G31: Other Degenerative Diseases of Nervous System |
| 30 | Bergè | 2011 | 21054282 | F20: Schizophrenia |
| 31 | Bergouignan L | 2009 | 19071222 | F32-F33: Major depressive disorder. single episode/recurrent |
| 32 | Berlingeri M | 2008 | 18413913 | G30: Alzheimer's Disease |
| 33 | Bernasconi N | 2004 | 15488421 | G40: Epilepsy and Recurrent Seizures |
| 34 | Bertsch K | 2013 | 23381548 | F60: Specific Personality Disorders |
| 35 | Beste C | 2008 | 17497629 | G10: Huntington's Disease |
| 36 | Beyer M K | 2007 | 17028119 | G20: Parkinson's Disease |
| 37 | Biundo R | 2011 | 21862438 | G20: Parkinson's Disease |
| 38 | Boccardi M | 2005 | 15585344 | G31: Other Degenerative Diseases of Nervous System |
| 39 | Bodini B | 2009 | 19172648 | G35: Multiple Sclerosis |
| 40 | Boghi A | 2011 | 21546219 | F50: Eating Disorders |
| 41 | Bonavita S | 2011 | 21239414 | G35: Multiple Sclerosis |
| 42 | Bonilha L | 2008 | 18164594 | F20: Schizophrenia |
| 43 | Bonilha L | 2004 | 15364683 | G40: Epilepsy and Recurrent Seizures |
| 44 | Borgwardt S J | 2010 | 20006324 | F20: Schizophrenia |
| 45 | Borroni B | 2008 | 18541800 | G31: Other Degenerative Diseases of Nervous System |
| 46 | Bose S K | 2009 | 19450953 | F20: Schizophrenia |
| 47 | Bouilleret V | 2008 | 18195263 | G40: Epilepsy and Recurrent Seizures |
| 48 | Boxer A L | 2006 | 16401739 | G23: Other degenerative diseases of basal ganglia |
| 49 | Boxer A L | 2003 | 12873851 | G30: Alzheimer's Disease |
| 50 | Boxer A L | 2003 | 12873851 | G31: Other Degenerative Diseases of Nervous System |
| 51 | Boxer A L | 2006 | 16401739 | G31: Other Degenerative Diseases of Nervous System |
| 52 | Bozzali M | 2006 | 16894107 | G30: Alzheimer's Disease |
| 53 | Bozzali M | 2006 | 16894107 | G31: Other Degenerative Diseases of Nervous System |
| 54 | Brambati S M | 2009 | 17604879 | G31: Other Degenerative Diseases of Nervous System |
| 55 | Brazdil M | 2009 | 18609565 | G40: Epilepsy and Recurrent Seizures |
| 56 | Brenneis C | 2004 | 14742598 | G23: Other degenerative diseases of basal ganglia |
| 57 | Brenneis C | 2004 | 15257132 | G30: Alzheimer's Disease |
| 58 | Brenneis C | 2004 | 15257132 | G31: Other Degenerative Diseases of Nervous System |
| 59 | Brenneis C | 2003 | 14534916 | G90: Disorders Autonomic Nervous System |
| 60 | Brenneis C | 2006 | 16161039 | G90: Disorders Autonomic Nervous System |
| 61 | Brieber S | 2007 | - | F84: Pervasive Developmental Disorders |
| 62 | Brieber S | 2007 | - | F90: Attention Deficit/Hyperactivity Disorder |
| 63 | Brunner R | 2010 | 19660555 | F60: Specific Personality Disorders |
| 64 | Burton E J | 2004 | 14749292 | G20: Parkinson's Disease |
| 65 | Burton E J | 2002 | 12377138 | G31: Other Degenerative Diseases of Nervous System |
| 66 | Cai Y | 2015 | 25502401 | F31: Bipolar Disorder |
| 67 | Cai Y | 2015 | 25502401 | F32-F33: Major depressive disorder. single episode/recurrent |
| 68 | Camicioli R | 2009 | 18573676 | G20: Parkinson's Disease |
| 69 | Canu E | 2010 | 21074899 | G30: Alzheimer's Disease |
| 70 | Carmona S | 2005 | 16129560 | F90: Attention Deficit/Hyperactivity Disorder |
| 71 | Caroli A | 2007 | 17990057 | G30: Alzheimer's Disease |
| 72 | Cascella N | 2010 | 20452187 | F20: Schizophrenia |
| 73 | Castro-Fornieles J | 2009 | 18486147 | F50: Eating Disorders |
| 74 | Castro-Manglano P D | 2011 | 22017223 | F20: Schizophrenia |
| 75 | Ceccarelli A | 2009 | 19172642 | G35: Multiple Sclerosis |
| 76 | Ceccarelli A | 2008 | 18501636 | G35: Multiple Sclerosis |
| 77 | Celle S | 2010 | 19768657 | G25: Other extrapyramidal and movement disorders |
| 78 | Ceresa A | 2013 | 23271221 | G35: Multiple Sclerosis |
| 79 | Chan C H | 2006 | 16499767 | G40: Epilepsy and Recurrent Seizures |
| 80 | Chaney A | 2014 | 23900024 | F32-F33: Major depressive disorder. single episode/recurrent |
| 81 | Chang C C | 2009 | 19486137 | G90: Disorders Autonomic Nervous System |
| 82 | Chang J L | 2005 | 16009889 | G12: Spinal Muscular Atrophy and Related Syndromes |
| 83 | Chao L L | 2012 | 22453299 | F43: Reaction to Severe Stress and Adjustment Disorders |
| 84 | Chen S | 2009 | 19538748 | F43: Reaction to Severe Stress and Adjustment Disorders |
| 85 | Chen S | 2006 | 16371250 | F43: Reaction to Severe Stress and Adjustment Disorders |
| 86 | Chen X | 2007 | 17464719 | F31: Bipolar Disorder |
| 87 | Chen Y | 2012 | 23155380 | F43: Reaction to Severe Stress and Adjustment Disorders |
| 88 | Cheng B | 2015 | 26347628 | F43: Reaction to Severe Stress and Adjustment Disorders |
| 89 | Cheng Y | 2010 | 20594947 | F32-F33: Major depressive disorder. single episode/recurrent |
| 90 | Cheng Y | 2011 | 21541322 | F84: Pervasive Developmental Disorders |
| 91 | Chetelat G | 2002 | 12395096 | G30: Alzheimer's Disease |
| 92 | Chetelat G | 2002 | 12395096 | G31: Other Degenerative Diseases of Nervous System |
| 93 | Chow E W | 2011 | 21362743 | F20: Schizophrenia |
| 94 | Chua S E | 2007 | 17098398 | F20: Schizophrenia |
| 95 | Compta Y | 2012 | 22595621 | G20: Parkinson's Disease |
| 96 | Cooke M A | 2008 | 18539438 | F20: Schizophrenia |
| 97 | Corbo V | 2005 | 16038682 | F43: Reaction to Severe Stress and Adjustment Disorders |
| 98 | Cordato N J | 2005 | 15843423 | G20: Parkinson's Disease |
| 99 | Cordato N J | 2005 | 15843423 | G23: Other degenerative diseases of basal ganglia |
| 100 | Cormack F | 2005 | 16006149 | G93: Other disorders of brain |
| 101 | Cosottini M | 2012 | 22226599 | G12: Spinal Muscular Atrophy and Related Syndromes |
| 102 | Craig M C | 2007 | 17766762 | F84: Pervasive Developmental Disorders |
| 103 | Critchley H D | 2003 | 12725766 | G90: Disorders Autonomic Nervous System |
| 104 | Cui L | 2011 | 21138758 | F20: Schizophrenia |
| 105 | Cui L | 2011 | 21138758 | F31: Bipolar Disorder |
| 106 | de Araujo-Filho G M | 2009 | 19303459 | G40: Epilepsy and Recurrent Seizures |
| 107 | de Oliveira-Souza R | 2008 | 18289882 | F60: Specific Personality Disorders |
| 108 | Delmaire C | 2007 | 17646630 | G24: Dystonia |
| 109 | Deng M Y | 2009 | 19641900 | F20: Schizophrenia |
| 110 | Di Paola M | 2007 | 17404777 | G30: Alzheimer's Disease |
| 111 | Dickstein D P | 2005 | 15997014 | F31: Bipolar Disorder |
| 112 | Doris A | 2004 | 15033185 | F31: Bipolar Disorder |
| 113 | Douaud G | 2007 | 17698497 | F20: Schizophrenia |
| 114 | Draganski B | 2003 | 14610125 | G24: Dystonia |
| 115 | Ebdrup B H | 2010 | 20184807 | F20: Schizophrenia |
| 116 | Eckart C | 2011 | 21118656 | F43: Reaction to Severe Stress and Adjustment Disorders |
| 117 | Ecker C | 2012 | 22310506 | F84: Pervasive Developmental Disorders |
| 118 | Ecker C | 2010 | 19683584 | F84: Pervasive Developmental Disorders |
| 119 | Egger K | 2008 | 19013058 | F32-F33: Major depressive disorder. single episode/recurrent |
| 120 | Ellis C M | 2001 | 11706094 | G12: Spinal Muscular Atrophy and Related Syndromes |
| 121 | Eshaghi A | 2014 | 3898881 | G35: Multiple Sclerosis |
| 122 | Etgen T | 2005 | 15670702 | G25: Other extrapyramidal and movement disorders |
| 123 | Euler M | 2009 | 19775870 | F20: Schizophrenia |
| 124 | Farrow T F D | 2005 | 15993858 | F31: Bipolar Disorder |
| 125 | Focke N K | 2011 | 21246668 | G20: Parkinson's Disease |
| 126 | Foong J | 2001 | 11335691 | F20: Schizophrenia |
| 127 | Friedrich H C | 2012 | 21967727 | F50: Eating Disorders |
| 128 | Frisoni G B | 2002 | 12438466 | G30: Alzheimer's Disease |
| 129 | Frodl T | 2008 | 18838632 | F32-F33: Major depressive disorder. single episode/recurrent |
| 130 | Gao W | 2013 | - | F31: Bipolar Disorder |
| 131 | Garcia-Marti G | 2008 | 17716795 | F20: Schizophrenia |
| 132 | Gaudio S | 2011 | 21081268 | F50: Eating Disorders |
| 133 | Gavazzi C | 2007 | 17882035 | G10: Huntington's Disease |
| 134 | Ghosh B C | 2012 | 22637582 | G23: Other degenerative diseases of basal ganglia |
| 135 | Gilbert A R | 2008 | 18342953 | F42: Obsessive Compulsive Disorder |
| 136 | Giordano A | 2013 | 23477861 | G23: Other degenerative diseases of basal ganglia |
| 137 | Giuliani N R | 2005 | 15721994 | F20: Schizophrenia |
| 138 | Gobbi C | 2014 | 23812284 | G35: Multiple Sclerosis |
| 139 | Gold B T | 2010 | 20063353 | G31: Other Degenerative Diseases of Nervous System |
| 140 | Gong Q | 2011 | 21134472 | F32-F33: Major depressive disorder. single episode/recurrent |
| 141 | Granert O | 2011 | 21705464 | G24: Dystonia |
| 142 | Gregory S | 2012 | 22566562 | F60: Specific Personality Disorders |
| 143 | Grieve S M | 2013 | 24273717 | F32-F33: Major depressive disorder. single episode/recurrent |
| 144 | Gross RG | 2010 | 20299856 | G31: Other Degenerative Diseases of Nervous System |
| 145 | Grosskreutz J | 2006 | 16638121 | G12: Spinal Muscular Atrophy and Related Syndromes |
| 146 | Guedj E | 2009 | 19224210 | G31: Other Degenerative Diseases of Nervous System |
| 147 | Guo W | 2014 | 24863419 | F32-F33: Major depressive disorder. single episode/recurrent |
| 148 | Guo X | 2010 | 19879920 | G30: Alzheimer's Disease |
| 149 | Ha T H | 2004 | 15664796 | F20: Schizophrenia |
| 150 | Ha T H | 2010 | 19429131 | F31: Bipolar Disorder |
| 151 | Hakamata Y | 2007 | 17923164 | F43: Reaction to Severe Stress and Adjustment Disorders |
| 152 | Haldane M | 2008 | 18308812 | F31: Bipolar Disorder |
| 153 | Hall A M | 2008 | 18631978 | G30: Alzheimer's Disease |
| 154 | Haller S | 2011 | 21284917 | F31: Bipolar Disorder |
| 155 | Hamalainen A | 2007 | 16997428 | G30: Alzheimer's Disease |
| 156 | Hamalainen A | 2007 | 16997428 | G31: Other Degenerative Diseases of Nervous System |
| 157 | Henley S M | 2009 | 19266143 | G10: Huntington's Disease |
| 158 | Herold R | 2009 | 19016669 | F20: Schizophrenia |
| 159 | Herringa R | 2012 | 23021615 | F43: Reaction to Severe Stress and Adjustment Disorders |
| 160 | Hirao K | 2008 | 18774263 | F20: Schizophrenia |
| 161 | Hirao K | 2006 | 16404228 | G30: Alzheimer's Disease |
| 162 | Honea R A | 2008 | 17689500 | F20: Schizophrenia |
| 163 | Honea R A | 2009 | 19812458 | G30: Alzheimer's Disease |
| 164 | Horn H | 2009 | 19182174 | F20: Schizophrenia |
| 165 | Horn H | 2010 | 20418073 | F20: Schizophrenia |
| 166 | Huey E D | 2009 | 19822784 | G31: Other Degenerative Diseases of Nervous System |
| 167 | Hulshoff Pol H E | 2001 | 11735840 | F20: Schizophrenia |
| 168 | Hulshoff Pol H E | 2004 | 14741639 | F20: Schizophrenia |
| 169 | Hulshoff Pol H E | 2006 | 16497519 | F20: Schizophrenia |
| 170 | Hwang J | 2010 | - | F32-F33: Major depressive disorder. single episode/recurrent |
| 171 | Hyde K L | 2010 | 19790171 | F84: Pervasive Developmental Disorders |
| 172 | Ille R | 2011 | 21406159 | G10: Huntington's Disease |
| 173 | Inkster B | 2011 | 20977527 | F32-F33: Major depressive disorder. single episode/recurrent |
| 174 | Ishii K | 2005 | 15800784 | G30: Alzheimer's Disease |
| 175 | Janssen J | 2008 | 18827723 | F20: Schizophrenia |
| 176 | Janssen J | 2008 | 18827723 | F28: Other psychotic disorder not due to a substance or known physiological condition |
| 177 | Janssen J | 2008 | 18827723 | F31: Bipolar Disorder |
| 178 | Jayakumar P N | 2005 | 15866362 | F20: Schizophrenia |
| 179 | Joos A | 2010 | 20400273 | F50: Eating Disorders |
| 180 | Kanda T | 2008 | 18661129 | G30: Alzheimer's Disease |
| 181 | Kanda T | 2008 | 18661129 | G31: Other Degenerative Diseases of Nervous System |
| 182 | Kasai K | 2008 | 17825801 | F43: Reaction to Severe Stress and Adjustment Disorders |
| 183 | Kasparek T | 2010 | 19777553 | F20: Schizophrenia |
| 184 | Kasparek T | 2007 | 17011096 | F20: Schizophrenia |
| 185 | Kasparek T | 2009 | 19647777 | F20: Schizophrenia |
| 186 | Kassubek J | 2005 | 15459079 | G10: Huntington's Disease |
| 187 | Kassubek J | 2004 | 14742591 | G10: Huntington's Disease |
| 188 | Kassubek J | 2007 | 17332050 | G12: Spinal Muscular Atrophy and Related Syndromes |
| 189 | Kato S | 2012 | 21850388 | G20: Parkinson's Disease |
| 190 | Kawachi T | 2006 | 16550383 | G30: Alzheimer's Disease |
| 191 | Kawada R | 2009 | 19625009 | F20: Schizophrenia |
| 192 | Kawasaki Y | 2004 | 15538599 | F20: Schizophrenia |
| 193 | Kawasaki Y | 2007 | 17045492 | F20: Schizophrenia |
| 194 | Ke X | 2008 | 18520994 | F84: Pervasive Developmental Disorders |
| 195 | Keller S S | 2007 | 17412561 | G40: Epilepsy and Recurrent Seizures |
| 196 | Keller S S | 2002 | 12438464 | G40: Epilepsy and Recurrent Seizures |
| 197 | Khaleeli Z | 2007 | 17566765 | G35: Multiple Sclerosis |
| 198 | Kim D | 2013 | 23769608 | F31: Bipolar Disorder |
| 199 | Kim E J | 2007 | 17615169 | G31: Other Degenerative Diseases of Nervous System |
| 200 | Kim J H | 2007 | 17689105 | G40: Epilepsy and Recurrent Seizures |
| 201 | Kim M J | 2008 | 18930633 | F32-F33: Major depressive disorder. single episode/recurrent |
| 202 | Kim S | 2011 | 21570296 | G30: Alzheimer's Disease |
| 203 | Kobel M | 2010 | 20702071 | F90: Attention Deficit/Hyperactivity Disorder |
| 204 | Koprivova J | 2009 | 19666084 | F42: Obsessive Compulsive Disorder |
| 205 | Kosaka H | 2010 | 20123027 | F84: Pervasive Developmental Disorders |
| 206 | Koskenkorva P | 2009 | 19704079 | G40: Epilepsy and Recurrent Seizures |
| 207 | Koutsouleris N | 2008 | 18054834 | F20: Schizophrenia |
| 208 | Kubicki M | 2002 | 12498745 | F20: Schizophrenia |
| 209 | Kubicki M | 2002 | 12498745 | F28: Other psychotic disorder not due to a substance or known physiological condition |
| 210 | Kurth F | 2011 | 21531390 | F84: Pervasive Developmental Disorders |
| 211 | Kwon H | 2004 | 15540637 | F84: Pervasive Developmental Disorders |
| 212 | Labate A | 2010 | 19780790 | G40: Epilepsy and Recurrent Seizures |
| 213 | Ladoucer C D | 2008 | 18356765 | F31: Bipolar Disorder |
| 214 | Lagarde J | 2013 | 24278277 | G23: Other degenerative diseases of basal ganglia |
| 215 | Lagarde J | 2013 | 24278277 | G31: Other Degenerative Diseases of Nervous System |
| 216 | Lai C H | 2015 | - | F32-F33: Major depressive disorder. single episode/recurrent |
| 217 | Lai C H | 2015 | - | F41: Other Anxiety Disorders |
| 218 | Lai C H | 2012 | 22386047 | F41: Other Anxiety Disorders |
| 219 | Lee H Y | 2011 | 21546094 | F32-F33: Major depressive disorder. single episode/recurrent |
| 220 | Leung K K | 2009 | 18945378 | F32-F33: Major depressive disorder. single episode/recurrent |
| 221 | Li C T | 2010 | 19931620 | F32-F33: Major depressive disorder. single episode/recurrent |
| 222 | Li L | 2006 | 16838824 | F43: Reaction to Severe Stress and Adjustment Disorders |
| 223 | Li M | 2011 | 21236649 | F31: Bipolar Disorder |
| 224 | Libon D J | 2009 | 19687454 | G31: Other Degenerative Diseases of Nervous System |
| 225 | Lin A | 2013 | 25206504 | G35: Multiple Sclerosis |
| 226 | Lin C H | 2013 | 23785322 | G20: Parkinson's Disease |
| 227 | Lin C H | 2013 | 23785322 | G25: Other extrapyramidal and movement disorders |
| 228 | Lin K | 2009 | 19570650 | G40: Epilepsy and Recurrent Seizures |
| 229 | Liu C H | 2014 | 24406440 | F32-F33: Major depressive disorder. single episode/recurrent |
| 230 | Liu M | 2011 | 22092238 | G40: Epilepsy and Recurrent Seizures |
| 231 | Lu C | 2010 | 19375076 | F80: Specific Developmental Disorders of Speech and Language |
| 232 | Ludolph A G | 2006 | 16648537 | F95: Tic Disorder |
| 233 | Lui S | 2009 | 18981063 | F20: Schizophrenia |
| 234 | Lyoo I K | 2004 | 15013835 | F31: Bipolar Disorder |
| 235 | Mainz V | 2012 | 22511729 | F50: Eating Disorders |
| 236 | Mak A K | 2009 | 19596037 | F32-F33: Major depressive disorder. single episode/recurrent |
| 237 | Mak E | 2013 | 24133286 | G20: Parkinson's Disease |
| 238 | Marcelis M | 2003 | 12694890 | F28: Other psychotic disorder not due to a substance or known physiological condition |
| 239 | Marti-Bonmati L | 2007 | 17641373 | F20: Schizophrenia |
| 240 | Massana G | 2003 | 12611840 | F41: Other Anxiety Disorders |
| 241 | Matsuda H | 2002 | 11884488 | G30: Alzheimer's Disease |
| 242 | Matsumoto R | 2010 | 20923432 | F42: Obsessive Compulsive Disorder |
| 243 | Matsunari I | 2007 | 18006622 | G30: Alzheimer's Disease |
| 244 | Mazere J | 2008 | 18191587 | G30: Alzheimer's Disease |
| 245 | McAlonan G M | 2005 | 15548557 | F84: Pervasive Developmental Disorders |
| 246 | McAlonan G M | 2002 | 12077008 | F84: Pervasive Developmental Disorders |
| 247 | McAlonan G M | 2008 | 18673405 | F84: Pervasive Developmental Disorders |
| 248 | McAlonan G M | 2007 | 17291727 | F90: Attention Deficit/Hyperactivity Disorder |
| 249 | McIntosh A M | 2004 | 15476683 | F20: Schizophrenia |
| 250 | McIntosh A M | 2004 | 15476683 | F31: Bipolar Disorder |
| 251 | McMillan A B | 2004 | 15325363 | G40: Epilepsy and Recurrent Seizures |
| 252 | Meda S A | 2008 | 18378428 | F20: Schizophrenia |
| 253 | Meisenzahl E M | 2008 | 18703313 | F20: Schizophrenia |
| 254 | Mengotti P | 2011 | 21146593 | F84: Pervasive Developmental Disorders |
| 255 | Meppelink A M | 2011 | 20922809 | G20: Parkinson's Disease |
| 256 | Mesaros S | 2008 | 18272867 | G35: Multiple Sclerosis |
| 257 | Mezzapesa D M | 2007 | 17296989 | G12: Spinal Muscular Atrophy and Related Syndromes |
| 258 | Miettinen P S | 2011 | 21692882 | G30: Alzheimer's Disease |
| 259 | Miettinen P S | 2011 | 21692882 | G31: Other Degenerative Diseases of Nervous System |
| 260 | Milham M P | 2005 | 15860335 | F41: Other Anxiety Disorders |
| 261 | Minnerop M | 2007 | 17512219 | G90: Disorders Autonomic Nervous System |
| 262 | Molina V | 2011 | 21188405 | F20: Schizophrenia |
| 263 | Molina V | 2010 | 20153145 | F20: Schizophrenia |
| 264 | Molina V | 2011 | 21188405 | F31: Bipolar Disorder |
| 265 | Moorhead T W | 2005 | 16085427 | F20: Schizophrenia |
| 266 | Morgen K | 2006 | 16360321 | G35: Multiple Sclerosis |
| 267 | Mueller S G | 2006 | 16686655 | G40: Epilepsy and Recurrent Seizures |
| 268 | Muhlau M | 2007 | 17024326 | G10: Huntington's Disease |
| 269 | Muhlau M | 2013 | 23462349 | G35: Multiple Sclerosis |
| 270 | Muller-Vahl K R | 2009 | 19435502 | F95: Tic Disorder |
| 271 | Na K S | 2013 | - | F41: Other Anxiety Disorders |
| 272 | Nagano-Saito A | 2005 | 15668417 | G20: Parkinson's Disease |
| 273 | Nardo D | 2013 | 23113800 | F43: Reaction to Severe Stress and Adjustment Disorders |
| 274 | Narita K | 2011 | 21115089 | F31: Bipolar Disorder |
| 275 | Neckelmann G | 2006 | - | F20: Schizophrenia |
| 276 | Nestor P J | 2003 | 12902311 | G31: Other Degenerative Diseases of Nervous System |
| 277 | Niedtfeldl I | 2013 | 23776553 | F60: Specific Personality Disorders |
| 278 | Nishio Y | 2010 | 20298422 | G20: Parkinson's Disease |
| 279 | Nugent A C | 2006 | 16256376 | F31: Bipolar Disorder |
| 280 | O'Daly O | 2007 | 17720459 | F20: Schizophrenia |
| 281 | O'Muircheartaigh J | 2011 | 21205693 | G40: Epilepsy and Recurrent Seizures |
| 282 | Obermann M | 2007 | 17443700 | G24: Dystonia |
| 283 | Ohnishi T | 2006 | 16330500 | F20: Schizophrenia |
| 284 | Ohnishi T | 2001 | 11673161 | G30: Alzheimer's Disease |
| 285 | Ortiz-Gil J | 2011 | 21727234 | F20: Schizophrenia |
| 286 | Overmeyer S | 2001 | 11722157 | F90: Attention Deficit/Hyperactivity Disorder |
| 287 | Padovani A | 2006 | 16306152 | G23: Other degenerative diseases of basal ganglia |
| 288 | Pail M | 2010 | 19817822 | G40: Epilepsy and Recurrent Seizures |
| 289 | Paillere-Martinot M L | 2001 | 11378311 | F20: Schizophrenia |
| 290 | Pantano P | 2011 | 20947646 | G24: Dystonia |
| 291 | Pardini M | 2009 | 19139305 | G31: Other Degenerative Diseases of Nervous System |
| 292 | Parisi L | 2014 | 24952616 | G35: Multiple Sclerosis |
| 293 | Peinemann A | 2005 | 16185716 | G10: Huntington's Disease |
| 294 | Pell G S | 2008 | 18042496 | G40: Epilepsy and Recurrent Seizures |
| 295 | Peng J | 2010 | 20466498 | F32-F33: Major depressive disorder. single episode/recurrent |
| 296 | Pennanen C | 2005 | 15607988 | G31: Other Degenerative Diseases of Nervous System |
| 297 | Pereira J B | 2009 | 19349926 | G20: Parkinson's Disease |
| 298 | Pereira J M | 2009 | 19433738 | G31: Other Degenerative Diseases of Nervous System |
| 299 | Pomarol-Clotet E | 2010 | 20065955 | F20: Schizophrenia |
| 300 | Prakash R S | 2010 | 19560443 | G35: Multiple Sclerosis |
| 301 | Preziosa P | 2016 | 26833969 | G35: Multiple Sclerosis |
| 302 | Price G | 2010 | 19632338 | F20: Schizophrenia |
| 303 | Pujol J | 2004 | 15237084 | F42: Obsessive Compulsive Disorder |
| 304 | Qiu L | 2011 | 21991357 | F20: Schizophrenia |
| 305 | Quattrone A | 2008 | 18653686 | G25: Other extrapyramidal and movement disorders |
| 306 | Rabinovici G D | 2007 | 18166607 | G30: Alzheimer's Disease |
| 307 | Rabinovici G D | 2007 | 18166607 | G31: Other Degenerative Diseases of Nervous System |
| 308 | Rami L | 2009 | 19259976 | G30: Alzheimer's Disease |
| 309 | Rami L | 2009 | 19259976 | G31: Other Degenerative Diseases of Nervous System |
| 310 | Ramirez-Ruiz B | 2007 | 17594330 | G20: Parkinson's Disease |
| 311 | Ranjeva J P | 2015 | 15661713 | G35: Multiple Sclerosis |
| 312 | Redlich R | 2014 | 25188810 | F31: Bipolar Disorder |
| 313 | Redlich R | 2014 | 25188810 | F32-F33: Major depressive disorder. single episode/recurrent |
| 314 | Remy F | 2005 | 15734360 | G30: Alzheimer's Disease |
| 315 | Riccitelli G | 2012 | 22422807 | G35: Multiple Sclerosis |
| 316 | Riederer F | 2008 | 18678824 | G40: Epilepsy and Recurrent Seizures |
| 317 | Ries M L | 2009 | 19701486 | F32-F33: Major depressive disorder. single episode/recurrent |
| 318 | Riva D | 2011 | 21700792 | F84: Pervasive Developmental Disorders |
| 319 | Rocca M A | 2014 | 24927473 | G35: Multiple Sclerosis |
| 320 | Rocha-Rego V | 2012 | 22952599 | F43: Reaction to Severe Stress and Adjustment Disorders |
| 321 | Rossi R | 2012 | 23146251 | F31: Bipolar Disorder |
| 322 | Rossi R | 2012 | 23146251 | F60: Specific Personality Disorders |
| 323 | Rusch N | 2004 | 15260365 | G40: Epilepsy and Recurrent Seizures |
| 324 | Salgado-Pineda P | 2003 | 12814586 | F20: Schizophrenia |
| 325 | Salgado-Pineda P | 2004 | 15006650 | F20: Schizophrenia |
| 326 | Salgado-Pineda P | 2011 | 21095105 | F20: Schizophrenia |
| 327 | Salmond C H | 2007 | 17710821 | F84: Pervasive Developmental Disorders |
| 328 | Salvadore G | 2011 | 21073959 | F32-F33: Major depressive disorder. single episode/recurrent |
| 329 | Sanchis-Segura C | 2016 | 27436479 | G35: Multiple Sclerosis |
| 330 | Santana M | 2010 | 20223639 | G40: Epilepsy and Recurrent Seizures |
| 331 | Saricicek A | 2015 | 26233321 | F31: Bipolar Disorder |
| 332 | Sasayama D | 2010 | 20546170 | F90: Attention Deficit/Hyperactivity Disorder |
| 333 | Saykin A J | 2006 | 16966547 | G31: Other Degenerative Diseases of Nervous System |
| 334 | Scheuerecker J | 2010 | 20569645 | F32-F33: Major depressive disorder. single episode/recurrent |
| 335 | Schiffer B | 2013 | 23015687 | F20: Schizophrenia |
| 336 | Schiffer B | 2007 | 16876824 | F65: Paraphilias |
| 337 | Schmidt-Wilcke T | 2009 | 19442751 | G31: Other Degenerative Diseases of Nervous System |
| 338 | Schuster C | 2012 | 21205677 | F20: Schizophrenia |
| 339 | Seeley W W | 2008 | 18268196 | G31: Other Degenerative Diseases of Nervous System |
| 340 | Senda J | 2011 | 21271792 | G12: Spinal Muscular Atrophy and Related Syndromes |
| 341 | Sepulcre J | 2006 | 16908748 | G35: Multiple Sclerosis |
| 342 | Serra-Blasco M | 2013 | 23620451 | F32-F33: Major depressive disorder. single episode/recurrent |
| 343 | Shad M U | 2012 | 22537357 | F32-F33: Major depressive disorder. single episode/recurrent |
| 344 | Shah P J | 1998 | 9828995 | F32-F33: Major depressive disorder. single episode/recurrent |
| 345 | Shapleske J | 2002 | 12427683 | F20: Schizophrenia |
| 346 | Shiino A | 2006 | 16904912 | G30: Alzheimer's Disease |
| 347 | Shiino A | 2006 | 16904912 | G31: Other Degenerative Diseases of Nervous System |
| 348 | Shin S | 2012 | 22933812 | G20: Parkinson's Disease |
| 349 | Sigmundsson T | 2001 | 11156806 | F20: Schizophrenia |
| 350 | Singh M K | 2012 | 3433284 | F31: Bipolar Disorder |
| 351 | Smesny S | 2010 | 20478385 | F20: Schizophrenia |
| 352 | Sobanski T | 2010 | 20056020 | F41: Other Anxiety Disorders |
| 353 | Soriano-Mas C | 2011 | 20875637 | F32-F33: Major depressive disorder. single episode/recurrent |
| 354 | Spano B | 2010 | 20007429 | G35: Multiple Sclerosis |
| 355 | Specht K | 2003 | 14568814 | G90: Disorders Autonomic Nervous System |
| 356 | Specht K | 2005 | 15734363 | G90: Disorders Autonomic Nervous System |
| 357 | Spencer M D | 2006 | 16996749 | F84: Pervasive Developmental Disorders |
| 358 | Stanfield A C | 2009 | 19267696 | F31: Bipolar Disorder |
| 359 | Stratmann M | 2014 | 25051163 | F32-F33: Major depressive disorder. single episode/recurrent |
| 360 | Suchan B | 2010 | 19729041 | F50: Eating Disorders |
| 361 | Sui S G | 2010 | - | F43: Reaction to Severe Stress and Adjustment Disorders |
| 362 | Summerfield C | 2005 | 15710857 | G20: Parkinson's Disease |
| 363 | Suzuki M | 2002 | 11955962 | F20: Schizophrenia |
| 364 | Szeszko P R | 2008 | 18413702 | F42: Obsessive Compulsive Disorder |
| 365 | Tae W S | 2006 | 16969045 | G40: Epilepsy and Recurrent Seizures |
| 366 | Tae W S | 2010 | 20046492 | G40: Epilepsy and Recurrent Seizures |
| 367 | Takahashi R | 2011 | 22187545 | G23: Other degenerative diseases of basal ganglia |
| 368 | Takahashi R | 2010 | 20634303 | G30: Alzheimer's Disease |
| 369 | Takahashi R | 2010 | 20634303 | G31: Other Degenerative Diseases of Nervous System |
| 370 | Taki Y | 2005 | 16150493 | F32-F33: Major depressive disorder. single episode/recurrent |
| 371 | Tang L R | 2014 | 25218414 | F31: Bipolar Disorder |
| 372 | Tavanti M | 2012 | 21710131 | F43: Reaction to Severe Stress and Adjustment Disorders |
| 373 | Tavazzi E | 2012 | 25228014 | G12: Spinal Muscular Atrophy and Related Syndromes |
| 374 | Tessitore A | 2012 | 22538070 | G20: Parkinson's Disease |
| 375 | Theberge J | 2007 | 17906243 | F20: Schizophrenia |
| 376 | Thivard L | 2007 | 17635981 | G12: Spinal Muscular Atrophy and Related Syndromes |
| 377 | Thomaes K | 2010 | 20673548 | F43: Reaction to Severe Stress and Adjustment Disorders |
| 378 | Tian L | 2011 | 22174900 | F20: Schizophrenia |
| 379 | Tiihonen J | 2008 | 18662866 | F60: Specific Personality Disorders |
| 380 | Tir M | 2009 | 19194988 | G20: Parkinson's Disease |
| 381 | Tir M | 2009 | 19194988 | G90: Disorders Autonomic Nervous System |
| 382 | Toal F | 2010 | 19891805 | F84: Pervasive Developmental Disorders |
| 383 | Togao O | 2010 | 20833001 | F42: Obsessive Compulsive Disorder |
| 384 | Tomelleri L | 2009 | 19717283 | F20: Schizophrenia |
| 385 | Tost H | 2010 | 19419772 | F31: Bipolar Disorder |
| 386 | Tregallas J R | 2007 | 17890058 | F20: Schizophrenia |
| 387 | Tzarouchi L C | 2010 | 19187475 | G90: Disorders Autonomic Nervous System |
| 388 | Uchida R R | 2008 | 18417322 | F41: Other Anxiety Disorders |
| 389 | Valente A A Jr | 2005 | 15978549 | F42: Obsessive Compulsive Disorder |
| 390 | van de Pavert S H | 2015 | 25926483 | G35: Multiple Sclerosis |
| 391 | van den Heuvel O A | 2009 | 18952675 | F42: Obsessive Compulsive Disorder |
| 392 | van Eijndhoven P | 2013 | 23929204 | F32-F33: Major depressive disorder. single episode/recurrent |
| 393 | van Tol M J | 2010 | 20921116 | F32-F33: Major depressive disorder. single episode/recurrent |
| 394 | van Tol M J | 2013 | 24176247 | F32-F33: Major depressive disorder. single episode/recurrent |
| 395 | van Tol M J | 2010 | 20921116 | F41: Other Anxiety Disorders |
| 396 | Venkatasubramanian G | 2008 | 19019637 | F20: Schizophrenia |
| 397 | Voets N L | 2008 | 18793730 | F20: Schizophrenia |
| 398 | Wagner G | 2008 | 18592043 | F32-F33: Major depressive disorder. single episode/recurrent |
| 399 | Wagner G | 2011 | 20832482 | F32-F33: Major depressive disorder. single episode/recurrent |
| 400 | Wang F | 2011 | 21666263 | F31: Bipolar Disorder |
| 401 | Wang J | 2007 | 1735333 | F90: Attention Deficit/Hyperactivity Disorder |
| 402 | Waragai M | 2009 | 19552926 | G30: Alzheimer's Disease |
| 403 | Watkins K E | 2002 | 11872605 | F80: Specific Developmental Disorders of Speech and Language |
| 404 | Watson D R | 2012 | 22056751 | F20: Schizophrenia |
| 405 | Watson D R | 2012 | 22056751 | F31: Bipolar Disorder |
| 406 | Wei W | 2016 | 26962820 | G40: Epilepsy and Recurrent Seizures |
| 407 | Whitford T J | 2006 | 16677830 | F20: Schizophrenia |
| 408 | Whitwell J L | 2013 | 23078273 | G23: Other degenerative diseases of basal ganglia |
| 409 | Whitwell J L | 2007 | 16797786 | G30: Alzheimer's Disease |
| 410 | Whitwell J L | 2005 | 16157747 | G31: Other Degenerative Diseases of Nervous System |
| 411 | Whitwell J L | 2007 | 16797786 | G31: Other Degenerative Diseases of Nervous System |
| 412 | Whitwell J L | 2007 | 17240166 | G31: Other Degenerative Diseases of Nervous System |
| 413 | Wilke M | 2001 | 11304078 | F20: Schizophrenia |
| 414 | Wilson S M | 2010 | 20542982 | G31: Other Degenerative Diseases of Nervous System |
| 415 | Woermann F G | 2000 | 10644781 | G40: Epilepsy and Recurrent Seizures |
| 416 | Wolf R C | 2008 | 18434103 | F20: Schizophrenia |
| 417 | Wolf R C | 2009 | 18172852 | G10: Huntington's Disease |
| 418 | Xie S | 2006 | 16801648 | G30: Alzheimer's Disease |
| 419 | Xu L | 2009 | 18266214 | F20: Schizophrenia |
| 420 | Yamada M | 2007 | 17240165 | F20: Schizophrenia |
| 421 | Yasuda C L | 2010 | 20350980 | G40: Epilepsy and Recurrent Seizures |
| 422 | Yoneyama E | 2003 | 14531753 | F60: Specific Personality Disorders |
| 423 | Yoo H K | 2005 | 16262646 | F41: Other Anxiety Disorders |
| 424 | Yoo S Y | 2008 | 18303194 | F42: Obsessive Compulsive Disorder |
| 425 | Yoshihara Y | 2008 | 19102744 | F20: Schizophrenia |
| 426 | Zahn R | 2005 | 16253483 | G30: Alzheimer's Disease |
| 427 | Zamboni G | 2008 | 18765649 | G31: Other Degenerative Diseases of Nervous System |
| 428 | Zhang J | 2011 | 21498053 | F43: Reaction to Severe Stress and Adjustment Disorders |
| 429 | Zhang T | 2009 | 19211150 | F32-F33: Major depressive disorder. single episode/recurrent |
| 430 | Zhang X | 2012 | 22129771 | F32-F33: Major depressive disorder. single episode/recurrent |
| 431 | Zhang X | 2016 | 28035997 | G35: Multiple Sclerosis |
| 432 | Zou K | 2010 | 19897176 | F32-F33: Major depressive disorder. single episode/recurrent |

**Table S4.** Selected articles for the meta-analysis of gray matter increases (VBM Query 2).

| **ID** | **First Author** | **Year** | **Medline** | **ICD-10 Code** |
| --- | --- | --- | --- | --- |
| 1 | Abell F | 1999 | 10501551 | F84: Pervasive Developmental Disorders |
| 2 | Adler C M | 2005 | 15922309 | F31: Bipolar Disorder |
| 3 | Adler C M | 2007 | 17027928 | F31: Bipolar Disorder |
| 4 | Amico F | 2011 | 20964952 | F32-F33: Major depressive disorder. single episode/recurrent |
| 5 | Antonini G | 2004 | 15489397 | G12: Spinal Muscular Atrophy and Related Syndromes |
| 6 | Antonova E | 2005 | 16039619 | F20: Schizophrenia |
| 7 | Arnone D | 2009 | - | F32-F33: Major depressive disorder. single episode/recurrent |
| 8 | Arnone D | 2013 | 23128153 | F32-F33: Major depressive disorder. single episode/recurrent |
| 9 | Asami T | 2009 | 19560907 | F41: Other Anxiety Disorders |
| 10 | Bassitt D P | 2007 | 16960651 | F20: Schizophrenia |
| 11 | Baxter L C | 2006 | 16914835 | G30: Alzheimer's Disease |
| 12 | Beal D S | 2007 | 17632278 | F80: Specific Developmental Disorders of Speech and Language |
| 13 | Betting L E | 2006 | 16702001 | G40: Epilepsy and Recurrent Seizures |
| 14 | Biederman S V | 2015 | 25809140 | F32-F33: Major depressive disorder. single episode/recurrent |
| 15 | Bonilha L | 2004 | 15364683 | G40: Epilepsy and Recurrent Seizures |
| 16 | Bonilha L | 2008 | 18362056 | F84: Pervasive Developmental Disorders |
| 17 | Bonilha L | 2007 | 17012334 | G40: Epilepsy and Recurrent Seizures |
| 18 | Brieber S | 2007 | - | F84: Pervasive Developmental Disorders |
| 19 | Brieber S | 2007 | - | F90: Attention Deficit/Hyperactivity Disorder |
| 20 | Brown G G | 2011 | 21924872 | F20: Schizophrenia |
| 21 | Brown G G | 2011 | 21924872 | F31: Bipolar Disorder |
| 22 | Butler C R | 2009 | 19073652 | G40: Epilepsy and Recurrent Seizures |
| 23 | Calderoni S | 2012 | 21896334 | F84: Pervasive Developmental Disorders |
| 24 | Calhoun V D | 2006 | 16108017 | F20: Schizophrenia |
| 25 | Carrion V G | 2009 | 19349151 | F43: Reaction to Severe Stress and Adjustment Disorders |
| 26 | Castro-Fornieles J | 2009 | 18486147 | F50: Eating Disorders |
| 27 | Celle S | 2010 | 19768657 | G25: Other extrapyramidal and movement disorders |
| 28 | Chan C H | 2006 | 16499767 | G40: Epilepsy and Recurrent Seizures |
| 29 | Chaney A | 2014 | 23900024 | F32-F33: Major depressive disorder. single episode/recurrent |
| 30 | Chen X | 2007 | 17464719 | F31: Bipolar Disorder |
| 31 | Chen Z | 2012 | 22119745 | F31: Bipolar Disorder |
| 32 | Cheng Y | 2011 | 21541322 | F84: Pervasive Developmental Disorders |
| 33 | Christian C J | 2008 | 18938065 | F42: Obsessive Compulsive Disorder |
| 34 | Cui L | 2011 | 21138758 | F20: Schizophrenia |
| 35 | Cui L | 2011 | 21138758 | F31: Bipolar Disorder |
| 36 | de Araujo-Filho G M | 2009 | 19303459 | G40: Epilepsy and Recurrent Seizures |
| 37 | de Castro-Manglano P | 2011 | 21316203 | F28: Other psychotic disorder not due to a substance or known physiological condition |
| 38 | Deng M Y | 2009 | 19641900 | F20: Schizophrenia |
| 39 | Ecker C | 2012 | 22310506 | F84: Pervasive Developmental Disorders |
| 40 | Ecker C | 2010 | 19683584 | F84: Pervasive Developmental Disorders |
| 41 | Egger K | 2007 | 17588241 | G24: Dystonia |
| 42 | Etgen T | 2005 | 15670702 | G25: Other extrapyramidal and movement disorders |
| 43 | Frangou S | 2012 | 3277296 | F31: Bipolar Disorder |
| 44 | Garraux G | 2006 | 16437578 | F95: Tic Disorder |
| 45 | Garraux G | 2004 | 15122716 | G24: Dystonia |
| 46 | Gee J | 2003 | 14697007 | G30: Alzheimer's Disease |
| 47 | Gilbert A R | 2008 | 18342953 | F42: Obsessive Compulsive Disorder |
| 48 | Giuliani N R | 2005 | 15721994 | F20: Schizophrenia |
| 49 | Gong Q | 2011 | 21134472 | F32-F33: Major depressive disorder. single episode/recurrent |
| 50 | Granert O | 2011 | 21705464 | G24: Dystonia |
| 51 | Grieve S M | 2013 | 24273717 | F32-F33: Major depressive disorder. single episode/recurrent |
| 52 | Grossman M | 2004 | 14761903 | G30: Alzheimer's Disease |
| 53 | Grossman M | 2004 | 14761903 | G31: Other Degenerative Diseases of Nervous System |
| 54 | Ha T H | 2010 | 19429131 | F31: Bipolar Disorder |
| 55 | Ha T H | 2004 | 15664796 | F20: Schizophrenia |
| 56 | Haldane M | 2008 | 18308812 | F31: Bipolar Disorder |
| 57 | Hamalainen A | 2007 | 17683950 | G31: Other Degenerative Diseases of Nervous System |
| 58 | Hendry J | 2006 | 16214373 | F84: Pervasive Developmental Disorders |
| 59 | Henley S M | 2009 | 19266143 | G10: Huntington's Disease |
| 60 | Honea R A | 2008 | 17689500 | F20: Schizophrenia |
| 61 | Hornyak M | 2007 | 17512782 | G25: Other extrapyramidal and movement disorders |
| 62 | Hulshoff Pol H E | 2001 | 11735840 | F20: Schizophrenia |
| 63 | Hwang J | 2010 | - | F32-F33: Major depressive disorder. single episode/recurrent |
| 64 | Hyde K L | 2010 | 19790171 | F84: Pervasive Developmental Disorders |
| 65 | Hyde T M | 2008 | 18669483 | F20: Schizophrenia |
| 66 | Kasparek T | 2010 | 19777553 | F20: Schizophrenia |
| 67 | Kawasaki Y | 2004 | 15538599 | F20: Schizophrenia |
| 68 | Ke X | 2008 | 18520994 | F84: Pervasive Developmental Disorders |
| 69 | Keller S S | 2007 | 17412561 | G40: Epilepsy and Recurrent Seizures |
| 70 | Keller S S | 2002 | 12438464 | G40: Epilepsy and Recurrent Seizures |
| 71 | Kempton M J | 2009 | 19726644 | F31: Bipolar Disorder |
| 72 | Kim J H | 2007 | 17689105 | G40: Epilepsy and Recurrent Seizures |
| 73 | Kim J J | 2001 | 11581113 | F42: Obsessive Compulsive Disorder |
| 74 | Kostic V S | 2010 | 20686125 | G20: Parkinson's Disease |
| 75 | Kozicky J M | 2013 | 23919287 | F31: Bipolar Disorder |
| 76 | Ladoucer C D | 2008 | 18356765 | F31: Bipolar Disorder |
| 77 | Lee S H | 2013 | 23474765 | G20: Parkinson's Disease |
| 78 | Leung K K | 2009 | 18945378 | F32-F33: Major depressive disorder. single episode/recurrent |
| 79 | Lin C H | 2013 | 23785322 | G20: Parkinson's Disease |
| 80 | Lin C H | 2013 | 23785322 | G25: Other extrapyramidal and movement disorders |
| 81 | Lin K | 2009 | 19570650 | G40: Epilepsy and Recurrent Seizures |
| 82 | Lu C | 2010 | 19375076 | F80: Specific Developmental Disorders of Speech and Language |
| 83 | Ludolph A G | 2006 | 16648537 | F95: Tic Disorder |
| 84 | Mallik S | 2015 | 4390521 | G35: Multiple Sclerosis |
| 85 | Marcelis M | 2003 | 12694890 | F28: Other psychotic disorder not due to a substance or known physiological condition |
| 86 | McDonald C | 2005 | 15863740 | F20: Schizophrenia |
| 87 | Mengotti P | 2011 | 21146593 | F84: Pervasive Developmental Disorders |
| 88 | Molina V | 2011 | 21188405 | F20: Schizophrenia |
| 89 | Moriya J | 2010 | 19854618 | F20: Schizophrenia |
| 90 | O'Daly O | 2007 | 17720459 | F20: Schizophrenia |
| 91 | Obermann M | 2007 | 17443700 | G24: Dystonia |
| 92 | Perico C A M | 2011 | 21320250 | F31: Bipolar Disorder |
| 93 | Perico C A M | 2011 | 21320250 | F32-F33: Major depressive disorder. single episode/recurrent |
| 94 | Prell T | 2013 | 24131497 | G24: Dystonia |
| 95 | Price G | 2010 | 19632338 | F20: Schizophrenia |
| 96 | Pujol J | 2004 | 15237084 | F42: Obsessive Compulsive Disorder |
| 97 | Qiu L | 2014 | 24713859 | F32-F33: Major depressive disorder. single episode/recurrent |
| 98 | Raji C A | 2009 | 19846828 | G30: Alzheimer's Disease |
| 99 | Ramirez-Ruiz B | 2007 | 17594330 | G20: Parkinson's Disease |
| 100 | Riederer F | 2008 | 18678824 | G40: Epilepsy and Recurrent Seizures |
| 101 | Rocca M A | 2015 | 26348234 | G35: Multiple Sclerosis |
| 102 | Rosen H J | 2002 | 11805245 | G31: Other Degenerative Diseases of Nervous System |
| 103 | Rusch N | 2004 | 15260365 | G40: Epilepsy and Recurrent Seizures |
| 104 | Salgado-Pineda P | 2003 | 12814586 | F20: Schizophrenia |
| 105 | Salmond C H | 2007 | 17710821 | F84: Pervasive Developmental Disorders |
| 106 | Saricicek A | 2015 | 26233321 | F31: Bipolar Disorder |
| 107 | Schafer A | 2010 | 20035881 | F50: Eating Disorders |
| 108 | Scheuerecker J | 2010 | 20569645 | F32-F33: Major depressive disorder. single episode/recurrent |
| 109 | Schiffer B | 2013 | 23015687 | F20: Schizophrenia |
| 110 | Schmitz N | 2006 | 16140278 | F84: Pervasive Developmental Disorders |
| 111 | Shapleske J | 2002 | 12427683 | F20: Schizophrenia |
| 112 | Smesny S | 2010 | 20478385 | F20: Schizophrenia |
| 113 | Suzuki M | 2002 | 11955962 | F20: Schizophrenia |
| 114 | Szeszko P R | 2008 | 18413702 | F42: Obsessive Compulsive Disorder |
| 115 | Tang L R | 2014 | 25218414 | F31: Bipolar Disorder |
| 116 | Tanskanen P | 2010 | 19015212 | F20: Schizophrenia |
| 117 | Tavazzi E | 2012 | 25228014 | G35: Multiple Sclerosis |
| 118 | Theberge J | 2007 | 17906243 | F20: Schizophrenia |
| 119 | Toal F | 2010 | 19891805 | F84: Pervasive Developmental Disorders |
| 120 | Truong W | 2013 | 24099630 | F32-F33: Major depressive disorder. single episode/recurrent |
| 121 | Uchida R R | 2008 | 18417322 | F41: Other Anxiety Disorders |
| 122 | Valente A A Jr | 2005 | 15978549 | F42: Obsessive Compulsive Disorder |
| 123 | van Eijndhoven P | 2013 | 23929204 | F32-F33: Major depressive disorder. single episode/recurrent |
| 124 | Waiter G D | 2004 | 15193590 | F84: Pervasive Developmental Disorders |
| 125 | Wang J | 2007 | 1735333 | F90: Attention Deficit/Hyperactivity Disorder |
| 126 | Watkins K E | 2002 | 11872605 | F80: Specific Developmental Disorders of Speech and Language |
| 127 | Watson D R | 2012 | 22056751 | F20: Schizophrenia |
| 128 | Watson D R | 2012 | 22056751 | F31: Bipolar Disorder |
| 129 | Wattendorf E | 2009 | 20007465 | G20: Parkinson's Disease |
| 130 | Whitford T J | 2006 | 16677830 | F20: Schizophrenia |
| 131 | Whitwell J L | 2004 | 16908994 | G31: Other Degenerative Diseases of Nervous System |
| 132 | Wilke M | 2001 | 11304078 | F20: Schizophrenia |
| 133 | Woermann F G | 1999 | 10545395 | G40: Epilepsy and Recurrent Seizures |
| 134 | Yasuda C L | 2010 | 20350980 | G40: Epilepsy and Recurrent Seizures |
| 135 | Yoo S Y | 2008 | 18303194 | F42: Obsessive Compulsive Disorder |

**Table S5.**  Sample characteristics for schizophrenia (F20: SCZ) and Alzheimer’s disease (G30: AD) datasets (BrainMap voxel-based morphometry sector).

| ***ICD-10 Code*** | ***Contrast*** | ***Articles*** | ***Experiments*** | ***Subjects*** |
| --- | --- | --- | --- | --- |
|  |  | ***(N)*** | ***(N)*** | ***(N)*** |
| F20: SCZ | Control > SCZ | 114 | 115 | 3807 |
|  | Control < SCZ | 28 | 33 | 1175 |
| G30: AD | Control > AD | 35 | 53 | 1194 |
|  | Control < AD | 4 | 22 | 114 |
|  | ***TOTAL*** | **181** | **230** | **6290** |

**Table S6.** Distribution of the functional experimental dataset.

Number of articles, experiments, and subjects for each selected paradigm class was reported.

| **Paradigm Class** | **Articles** | **Experiments** | **Subjects** |
| --- | --- | --- | --- |
|  | **(N)** | **(N)** | **(N)** |
| Finger tapping/button press | 193 | 1177 | 7011 |
| Reward | 106 | 728 | 3432 |
| Semantic monitor/discrimination | 109 | 616 | 2960 |
| Face monitoring/discrimination | 97 | 573 | 2755 |
| Emotion induction | 81 | 462 | 2790 |
| Passive viewing | 79 | 450 | 2186 |
| Cued Explicit Recognition/recall | 65 | 382 | 1791 |
| Visuospatial attention | 70 | 353 | 1693 |
| Pain monitor/discrimination | 63 | 325 | 1439 |
| N-back | 64 | 291 | 2609 |
| Film viewing | 42 | 266 | 1199 |
| Dealyed match to sample | 48 | 259 | 1397 |
| Theory of mind | 38 | 253 | 1197 |
| Go/No go | 50 | 246 | 1841 |
| Encoding | 48 | 236 | 1489 |
| Passive listening | 48 | 234 | 1153 |
| Counting/calculation | 44 | 232 | 1116 |
| Visual object identification | 33 | 229 | 853 |
| Affective pictures | 33 | 225 | 1326 |
| Phonological discrimination | 38 | 209 | 857 |
| Reasoning/problem solving | 36 | 206 | 1318 |
| Flexion/extension | 34 | 199 | 790 |
| Reading (covert) | 37 | 194 | 872 |
| Gambling | 22 | 159 | 702 |
| Music comprehension | 19 | 155 | 510 |
| Word generation (covert) | 35 | 155 | 779 |
| Pitch monitor/discrimination | 22 | 142 | 554 |
| Word generation (overt) | 28 | 142 | 782 |
| Olfactory monitoring/discrimination | 22 | 141 | 557 |
| Paired associate recall | 27 | 138 | 699 |
| Tactile monitor/discrimination | 28 | 134 | 580 |
| Reading (overt) | 26 | 131 | 595 |
| Vibrotactile monitor/discrimination | 8 | 130 | 169 |
| Imagined objects/scenes | 25 | 128 | 742 |
| Orthographic discrimination | 24 | 124 | 653 |
| Saccades | 31 | 123 | 574 |
| Sexual arousal/gratification | 17 | 122 | 609 |
| Naming (overt) | 20 | 120 | 467 |
| Tone monitor/discrimination | 27 | 119 | 643 |
| Episodic recall | 22 | 117 | 554 |
| Mental rotation | 19 | 117 | 493 |
| Task switching | 22 | 117 | 644 |
| Taste | 18 | 113 | 623 |
| Stroop - color | 30 | 110 | 1049 |
| Visual pursuit/tracker | 19 | 95 | 390 |
| Imagined movement | 20 | 94 | 522 |
| Chewing/swallowing | 15 | 90 | 397 |
| Classical conditioning | 17 | 90 | 377 |
| Meditation | 11 | 85 | 433 |
| Recitation/repetition (overt) | 19 | 78 | 408 |
| Music production | 17 | 77 | 407 |
| Naming (covert) | 14 | 73 | 288 |
| Sequence recall/learning | 14 | 65 | 334 |
| Affective words | 8 | 62 | 398 |
| Wisconsin card sorting test | 12 | 57 | 352 |
| Deception | 12 | 52 | 340 |
| Oddball discrimination | 12 | 48 | 376 |
| Grasping | 9 | 45 | 146 |
| Delay discounting | 5 | 42 | 262 |
| Drawing | 4 | 39 | 77 |
| Flanker | 10 | 37 | 266 |
| Rest | 11 | 36 | 386 |
| Transcranical Magnetic Stimulation | 8 | 35 | 129 |
| Pointing | 8 | 34 | 113 |
| Acupuncture | 7 | 33 | 193 |
| Figurative language | 6 | 33 | 160 |
| Flashing checkerboard | 3 | 33 | 185 |
| Object manipulation/discrimination | 4 | 32 | 58 |
| Anti Saccades | 8 | 31 | 202 |
| Recitation/repetition (covert) | 10 | 31 | 212 |
| Fixation | 9 | 30 | 185 |
| Competition/cooperation | 3 | 29 | 116 |
| Hunger/satiety | 6 | 26 | 134 |
| Isometric force | 5 | 26 | 104 |
| Lexical decision | 6 | 26 | 165 |
| Micturition | 6 | 25 | 128 |
| Divided auditory attention | 4 | 24 | 98 |
| Magnitude comparison (symbolic) | 5 | 24 | 126 |
| Self reflection | 2 | 24 | 87 |
| Stroop - emotional | 8 | 23 | 328 |
| Syntactic discrimination | 8 | 23 | 212 |
| HyperCapnia /air Hunger | 7 | 22 | 104 |
| Pursuit rotor/manual tracking | 4 | 21 | 75 |
| Magnitude comparison (phisical size) | 3 | 20 | 97 |
| Video games | 4 | 20 | 111 |
| Emotional body language perception | 2 | 17 | 51 |
| Driving | 3 | 15 | 69 |
| Estimation | 4 | 15 | 120 |
| Magnitude comparison (distance) | 1 | 15 | 38 |
| Tower of London | 4 | 15 | 84 |
| Multi tasking | 3 | 14 | 82 |
| Stroop - counting | 4 | 14 | 105 |
| Stroop - other | 3 | 14 | 103 |
| Word Stem completion (overt) | 4 | 14 | 103 |
| Magnitude comparison (luminance) | 2 | 13 | 37 |
| Magnitude comparison (numerical) | 2 | 13 | 37 |
| Word imageability | 3 | 13 | 84 |
| Free list word record | 4 | 11 | 69 |
| Thirst induction | 3 | 11 | 67 |
| Visual Motion | 2 | 11 | 55 |
| Writing | 3 | 11 | 83 |
| Stroop - spatial | 2 | 10 | 36 |
| Hand-Eye Coordination | 2 | 7 | 44 |
| Motor learning | 1 | 7 | 15 |
| Sleep | 3 | 7 | 77 |
| Trauma recall | 2 | 7 | 68 |
| Vestibular Stimulation | 3 | 7 | 54 |
| Induced panic | 2 | 6 | 62 |
| Word stem completion (covert) | 2 | 6 | 64 |
| Fluency induction | 1 | 3 | 12 |
| **TOTAL** | **2376** | **13148** | **68152** |
